# Maize Biochemical Defences against a Rootworm Were Mediated by Domestication, Spread, and Breeding

**DOI:** 10.1101/2020.06.03.132795

**Authors:** Ana A. Fontes-Puebla, Eli J. Borrego, Michael V. Kolomiets, Julio S. Bernal

## Abstract

Plant physiological processes generally are regulated by phytohormones, including plant biochemical responses to herbivory. Here, we addressed whether a suite of maize (*Zea mays mays*) phytohormones, including some precursor and derivative metabolites, relevant to herbivory defence were mediated by the crop’s domestication, northward spread, and modern breeding. For this, we compared phytohormone and metabolite levels among four plant types representing the evolutionary and agronomic transitions from maize’s wild ancestor, Balsas teosinte (*Zea mays parviglumis*), to Mexican and US maize landraces, and to highly-bred US maize cultivars, as affected by root herbivory by Western corn rootworm (*Diabrotica virgifera virgifera*). Following ecological-evolutionary hypotheses, we expected to find changes in: (i) maize defence strategy, from reliance on induced to constitutive defences; (ii) levels of phytohormones relevant to herbivore resistance consistent with gradual weakening of defences, and; (iii) levels of a phytohormone relevant to herbivory tolerance because it positively affects plant growth. We found that with its domestication, maize seemed to have transitioned from reliance on induced defences in Balsas teosinte to reliance on constitutive defences in maize. Also, we found that while one subset of phytohormones relevant to herbivory was suppressed (13-oxylipins), another was enhanced (9-oxylipins) with domestication, and both subsets were variably affected by spread and breeding. Finally, an auxin phytohormone directly linked to growth (indole-3-acetic acid), increased significantly with domestication, and seemingly with spread and breeding. We concluded that rootworm defences in maize were mediated by domestication and ensuing processes, such as spread and breeding, and argued that agricultural intensification mediated maize defence evolution in parallel with modern breeding.

## INTRODUCTION

Phytohormones, including precursor and derivative metabolites, regulate a multitude of plant processes, including biochemical defences against herbivores and pathogens, responses to biotic and abiotic stresses, and plant growth [1, 2]. For example, jasmonic acid (JA) has multiple functions in plants, among which are initiating defence responses against insects and pathogens, and inducing trichome development, among others [3-5]. Similarly, salicylic acid (SA) has been associated with seed germination, cell growth, stomatal closure, responses to abiotic stressors, and defence against biotrophic and hemibiotrophic pathogens, among other processes [6]. Importantly, cross-talk may occur between phytohormones, e.g., antagonism between JA- and SA-initiated responses to herbivores and pathogens, respectively [7, 8].

Phytohormones, including precursors and derivatives, are grouped in classes that include auxins (e.g., indole-3-acetic acid or IAA), jasmonates (e.g., JA), salicylates (e.g., SA), and others, and are produced in a variety of metabolic pathways. Specifically, jasmonates are produced by the lipoxygenase (LOX) pathway, and salicylates are derived from the chorismate pathway, whereas IAA is mainly synthesized from the amino acid tryptophan [1, 9-13]. While jasmonates and salicylates play well-known roles in biochemical herbivore defences, IAA may indirectly mediate herbivore defences by regulating plant growth and inducing a subset of jasmonate-related defences [6, 14, 15]. Production levels of phytohormones, and precursor and derivative metabolites vary within and between plant species, and can be shaped by a variety of selective pressures, including natural and artificial selection through herbivory, disease, crop domestication, systematic breeding, and others [16-18].

Jasmonates are representatives of an approximately 700-strong group of lipid-derived metabolites known as oxylipins. More specifically, jasmonates are members of large group of oxylipins produced by the initial oxidation at carbon position 13th of polyunsaturated fatty acids by 13-LOX isozymes, and thus are classified as 13-oxylipins [19, 20]. Another large class of biologically active metabolites are produced by 9-LOXs, and classified as 9-oxylipins, contain molecules with either signalling or direct insecticidal activities [19, 21-23].

Plant defence responses to aboveground herbivory have been studied extensively [14, 24, 25], while responses to belowground herbivory are poorly understood [26-29]. Nevertheless, some reports suggest that belowground herbivory and mechanical wounding trigger plant responses similar to those triggered in aboveground tissues [30, 31]. Generally, phytohormones and related metabolites act as signals responsible for modulating gene transcription leading to production of defensive compounds [30, 32-34]. Furthermore, phytohormones may travel to distant and undamaged tissues through plant vascular systems, preparing those tissues for potential attack by pathogens or insects [35].

Plants under herbivore attack allocate resources towards defence, which reduces the availability of resources for growth and reproduction. Thus, the Optimal Defence Hypothesis postulates that there is a cost for defence in plants, whether constitutive or induced [36]. However, how plants invest resources in herbivore defence seems to depend on their genetics, the availability of resources, and pressure from other stresses, among other variables [36-39]. Accordingly, crop domestication may affect how plants invest in herbivory defence by re-directing resources to productivity, including yield, rather than defence [38, 40]. Additionally, plant spread to new environments exposes them to novel herbivores or allows them to escape endemic herbivores, so may mediate the evolution of tolerance and resistance [36, 37]. Artificial selection and breeding in crop species have historically enhanced productivity (i.e., yield) and other human-interest traits over resistance to herbivores and pathogens, so may also mediate the evolution of tolerance and resistance [41-44]. Moreover, in contexts in which availability of resources is high, e.g., in agricultural compared to wild habitats, selection may favour fast growth and tolerance to herbivores and pathogens, rather than resistance [45]. Such may be the case for maize (*Zea mays mays* L.), which was cultivated in increasingly favourable contexts, and underwent successive bouts of natural and artificial selection, as it was domesticated, spread in the Americas and became a dietary staple, and, most recently, whilst subjected to systematic breeding for enhanced yield under intensive agriculture [17, 41, 46-49]. Indeed, prior studies showed that these processes significantly shaped how maize responds to herbivory [47, 50, 51].

In addition to domestication, spread, and breeding, previous studies suggested that herbivory pressure, resource availability, and agricultural intensification may mediate the evolution of maize defences against herbivores, including Western corn rootworm (WCR) (*Diabrotica virgifera virgifera* LeConte) [50, 51]. As for other crops and herbivores, herbivory pressure by WCR likely contributed to shaping maize defensive strategies, whether based on resistance or tolerance, after the crop spread from central Mexico to North America [37, 51-53]. Maize resistance (e.g., antibiosis) and tolerance (e.g., compensatory growth) to WCR depend on the triggering of signalling cues, particularly phytohormones. Particularly, resistance may imply synthesis of secondary metabolites triggered by changes in phytohormone levels, whereas tolerance may depend in part on synthesis of growth-related phytohormones [6, 12, 20]. Thus, constitutive and herbivore-induced phytohormone profiles relevant to maize resistance and tolerance to WCR, and other herbivores, may have been mediated by the crop’s domestication, spread, and systematic breeding, as well as by variable herbivory pressure, resource availability, and agricultural intensification [50, 51, 54].

In this study, we tested whether the profiles of constitutive and induced maize phytohormones, including precursor and derivative metabolites, relevant to defence were mediated by the crop’s domestication, spread, and breeding. Specifically, we compared the profiles of those phytohormones, precursors, and derivatives (hereafter “metabolites”) among four plant types representing the evolutionary and agronomic transitions from maize’s wild ancestor to highly-bred maize cultivars, viz.: Balsas teosinte (*Zea mays* L. spp. *parviglumis* Iltis and Doebley), Mexican maize landraces, US maize landraces, and US maize breeding lines. Each plant type was represented by three plant accessions to better capture the genetic diversity within each type. Domestication effects were assessed by comparing constitutive and induced metabolite profiles and levels between Balsas teosinte and Mexican maize landraces; of northward spread by comparing between Mexican landraces and US landraces, and; of breeding by comparing between US landraces and US inbred lines. Overall, from Balsas teosinte to US inbred maize lines we expected to find decreasing levels of metabolites positively related to resistance, and increasing levels of metabolites positively related to plant growth. Specifically, we expected to find: (i) a shift from reliance on induced to constitutive herbivore defence with domestication and cultivation in increasingly rich environments; (ii) a decline in levels of putative insecticidal metabolites of the LOX pathway, and; (iii) an increase in the growth-promoting phytohormone, IAA [37, 38, 40, 54]. We discussed our results in the contexts of those expectations, as well as in reference to prior results concerning the evolution of maize defence against WCR, as mediated by artificial and natural selection, geographical spread, and systematic breeding [51].

## MATERIALS AND METHODS

### Plants and Insects

Assays included four plant types spanning the evolution of maize from its domestication from its wild ancestor to its subsequent spread and systematic breeding in USA: (i) Balsas teosinte (immediate ancestor of maize); (ii) Mexican landraces (descendants of Balsas teosinte), which served to assess any domestication effects; (iii) US landraces (descendants of Mexican landraces), used to assess any effects of northward spread, and; (iv) US inbred lines (derived from US landraces), used for assessing any effects of systematic breeding [42, 55-59]. Each plant type included three accessions: “El Cuyotomate,” “Talpitita,” and “El Rodeo” for Balsas teosinte; Palomero Toluqueño, Chalqueño, and Cacahuacintle for Mexican landraces; Lancaster Sure Crop, Reid Yellow Dent, and Gourdseed for US landraces, and; Mo17, B73, and W438 for US inbred lines (Table 1) [51]. Hereafter, “BTEO” refers to Balsas teosinte, “MXLR” to Mexican landraces, “USLR” to US landraces, and “USIL” to US inbred lines.

**Table 1.** Plant types and accessions, and the geographic origin and reference number of each, where applicable. From top to bottom, plant types are ordered from most ancestral to most derived, to reflect the chronological sequence of the domestication, spread and breeding processes.

| PLANT TYPE/ACCESSION | ORIGIN | REFERENCE<sup>4</sup> |
| --- | --- | --- |
| **Balsas teosintes<sup>1</sup>** |  |  |
|  | *Jalisco state, Mexico:* |  |
| El Cuyotomate | Ejutla, Ejutla (19°58'N, 104°04'W) | — |
| Talpitita | Talpitita, Villa Purificación (19°42'N, 104°48'W) | — |
| El Rodeo | El Rodeo, Tolimán (19°33'N, 104°03'W) | — |
| **Mexican landraces<sup>2</sup>** |  |  |
|  | *Mexico state, Mexico:* |  |
| Palomero Toluqueño | Toluca Valley, Toluca | NSL 2824 |
| Chalqueño | San Mateo Atenco, San Mateo Atenco | PI 629215 |
| Cacahuacintle | Toluca Valley, Toluca | NSL 2823 |
| **US landraces<sup>2</sup>** |  |  |
|  | *United States:* |  |
| Lancaster Sure Crop | Ohio | PI 280061 |
| Reid Yellow Dent | Indiana | PI 213698 |
| Gourdseed | Ennis, Texas | PI 414179 |
| **US inbred lines** |  |  |
|  | *United States:* |  |
| Mo17<sup>2</sup> | Missouri | PI 558532 |
| B73<sup>3</sup> | Iowa | PI 550473 |
| W438<sup>3</sup> | Wisconsin | AMES 29447 |

<sup>1</sup>Collected by JSB; <sup>2</sup>Provided by USDA, ARS Germplasm Resources Information Network (GRIN);
<sup>3</sup>Provided by M. V. Kolomiets, Texas A&M University, College Station; <sup>4</sup>USDA, ARS GRIN reference number.

Seeds were germinated in Petri dishes (150 × 15 mm) within moistened, absorbent paper towels for 5 d (BTEO, USIL) or 4 d (MXLR, USLR), without prior seed surface sterilization. Teosinte seeds were removed from their fruitcases using a nail clipper before germinating. After germinating, each seedling was transplanted to a cone-tainer (4 × 25 cm, diam × length) (Stuewe & Sons, Tangent, OR, USA), modified with chiffon mesh to prevent escape of WCR larvae through its drainage holes, and allowed to grow for an additional 10 d. Growing conditions were 25 ± 2 °C, 50% RH, 12:12 photoperiod (L:D), and as-needed watering. Commercial potting soil (Baccto® Premium, Michigan Peat Co., Houston, TX, USA) was used in all assays, which was sifted through a 60-mesh sieve before transplanting seeds to facilitate subsequent root harvest (see below).

WCR eggs, of the diapause strain, and provided by USDA-ARS-North Central Agricultural Research Laboratory (Brookings, SD, USA) were used in all assays. These were incubated in Petri dishes (150 × 15 mm) at 25 ± 2 °C, ∼ 80% RH for 12 ± 1 d on moistened absorbent paper. Neonate first-instar larvae (< 24 h after hatching) were collected from the Petri dishes and used in all assays.

### Constitutive, total induced, and net induced metabolites

Constitutive and induced levels of metabolites (i.e., phytohormones, including precursors and derivatives), were assessed in maize seedlings free of WCR herbivory or exposed to WCR herbivory, respectively. The metabolites were selected because of their relevance to maize defence responses to insects, and included: 12-oxophytodienoic acid (hereafter 12-OPDA), jasmonic acid (JA), jasmonyl-isoleucine (JA-Ile), 12-carboxy-jasmonyl-isoleucine (12-COOH-JA-Ile), 10-oxo-phytoenoic acid (10-OPEA), 10-oxo-phytodienoic acid (10-OPDA), DA^0^-4:0, a 10-OPEA derivative with 4 carbons in the carboxylic side chain (DA4), Azelaic acid (AZA), coumaric acid (COU), benzoic acid (BNZ), salicylic acid (SA), and indole-3-acetic acid (IAA) (Figure 1). The selected metabolites occur in three biochemical pathways: lipoxygenase, phenylpropanoid, and tryptophan (Figure 1). The lipoxygenase (LOX) pathway, with two branches, 13-LOX-derived oxylipins (13-oxylipins) and 9-LOX-derived oxylipins (9-oxylipins), includes 12-OPDA, JA, JA-Ile, 12COOH-JA-Ile, 10-OPEA, 10-OPDA, DA4, AZA, and is broadly relevant to insect and pathogen defence [13, 19, 20, 22, 60]. The phenylpropanoid pathway (PAL), which includes COU, BNZ, and SA, is broadly relevant to pathogen defence [6, 10]. Finally, the auxin biosynthesis pathway (AUX) derived from the tryptophan pathway is broadly relevant to plant growth [12, 61].

**Figure 1.**
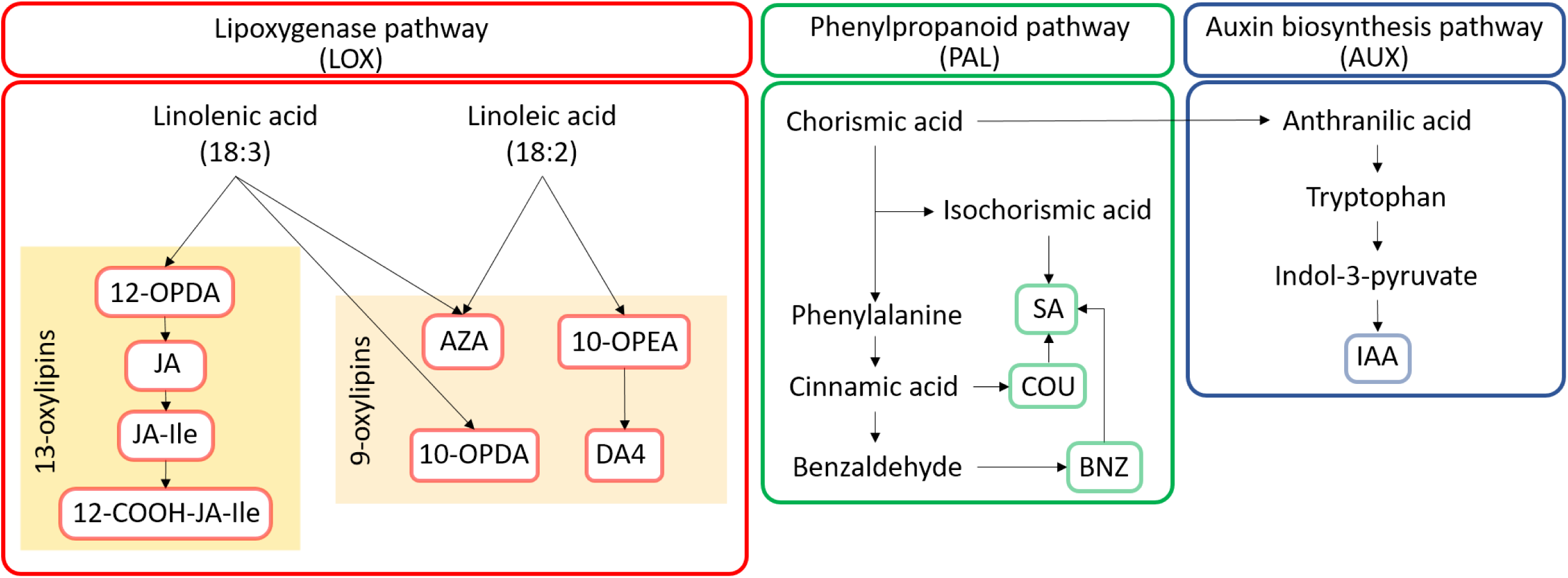
Place of the 12 defence-related metabolites measured in this study within three metabolic pathways relevant to plant defence and growth; pathways are presented in abbreviated form. The pathways and metabolites are indicated by their enclosure within distinctively-coloured rectangles. The lipoxygenase pathway (LOX) (red rectangles) includes 13-oxylipin and 9-oxylipin branches, with derivatives oxidized in the 13- and 9-carbon, respectively; it includes 12-oxo-phytodienoic acid (12-OPDA), jasmonic acid (JA), jasmonyl-isoleucine (JA-Ile), and 12-carboxy-jasmonyl-isoleucine (12-COOH-JA-Ile) within the 13-oxylipins branch, and 10-oxo-phytodienoic acid (10-OPDA), 10-oxo-phytoenoic acid (10-OPEA), DA4, and azelaic acid (AZA) within the 9-oxylipins branch. The phenylpropanoid pathway (PAL) (green rectangles) includes salicylic acid (SA), benzoic acid (BNZ), and coumaric acid (COU). Finally, the auxin biosynthesis pathway (AUX) (blue rectangles) includes only the Indole-3-acetic acid (IAA).

“Constitutive” metabolites, were assessed in seedlings free of WCR herbivory; “total induced” metabolites were assessed in seedlings exposed to WCR herbivory for 8, 24, or 48 h, and; “net induced” metabolites were estimated as the difference between total induced and constitutive metabolites, i.e., net induced metabolites = total induced metabolites – constitutive metabolites. For statistical analyses and interpretation, metabolite measurements at 8, 24, and 48 h were pooled and used as measures of total induced (and net induced) levels because the levels of individual metabolites peaked at varying exposure intervals across plant types and accessions (see below, *Statistical analyses*).

To measure metabolite levels, 10 neonate WCR larvae were placed on the soil surface in individual cone-tainers holding a ∼15 d-old seedling [62], and allowed to burrow and feed for intervals of 8, 24, or 48 h; in parallel, one set of seedlings (of size and number of leaves similar to those of seedlings exposed to larvae) was not exposed to WCR larvae. The assay included three biological replicates for each exposure interval, as well as for non-exposed seedlings, per each plant accession, i.e., nine biological replicates per plant type; each biological replicate consisted of four seedlings. After the exposure intervals, seedling roots were carefully cleansed of soil by rinsing with running water, after which they were excised, flash-frozen, and kept at −80 °C until ground; seedlings not exposed to WCR were similarly processed at 0 h, i.e., at the time WCR larvae were placed in exposed seedlings. A mortar and pestle were used to grind root tissues in liquid nitrogen. The ground tissue samples were kept at −80 °C until metabolite extraction [4, 23].

### Metabolite extraction and quantification

For extraction of metabolites in fresh tissue, a 104.3 ± 0.651 mg portion of ground tissue from each plant accession was mixed with 500 µL of alcohol-based phytohormone extraction buffer (1-propanol/water/HCl [2:1:0.002 v/v/v]) containing 5 µM of isotopically-labelled internal standards: d-ABA ([2H6](+)-cis, trans-abscisic acid; Olchemlm cat# 0342721), d-ACC (1-Aminocyclopropane-2,2,3,3-d4-carboxylic acid; Sigma cat#736260), d-IAA([2H5] indole-3-acetic acid; Olchemlm cat# 0311531), d-JA (2,4,4-d3; acetyl-2,2-d2 jasmonic acid; CDN Isotopes cat# D-6936), and d-SA (d6-salicylic acid; Sigma cat#616796), with further 30 min agitation at 4 °C in darkness; 500 µL of dichloromethane were added and agitated for another 30 min at 4 °C followed by centrifugation (13,000 g for 5 min) at 4 °C in darkness. The supernatant was removed and the remaining organic solvent was evaporated under N_2_ (g) flow. The pellet was re-solubilized in 150 µL of MeOH, shaken for 1 min and centrifuged (14,000 g for 2 min). The supernatant (90 µL) was analysed by liquid chromatography-mass spectrometry (API 3200, QqQ; Sciex, Framingham, MA) ran in negative ESI mode. The column used was an Ascentis Express 30 × 2.1 mm 2.7 µm solid core column following the mobile phases and elution conditions as described by [23].

### Statistical analyses

Preliminary analyses –two-way ANOVA including the interaction “time × plant type” with metabolite levels as dependent variables, and time (8, 24, and 48 h) and plant type as independent variables– showed that the levels of the 12 metabolites peaked at different exposure intervals (8 h, 24 h, 48 h) across the 12 plant accessions, so that inclusion of a single exposure interval in final analyses would not adequately represent the responses for all accessions and plant types. Consequently, statistical analyses considered herbivory by WCR as an independent variable with two levels: without herbivory (= not exposed to WCR, i.e., constitutive metabolite levels), and with herbivory (= exposed to WCR, independently of the duration of exposure, i.e., total induced metabolite levels).

Independent multivariate analyses of variance (MANOVA) were applied to evaluate whether “constitutive,” “total induced,” and “net induced” metabolite levels were affected by domestication, spread, and breeding. Constitutive and total induced metabolite levels were measured, respectively, as per-replicate levels in seedlings without exposure or with exposure to WCR; net induced levels were calculated as the per-replicate total induced level minus the corresponding per-accession, average constitutive level. The independent variables were “plant type” (BTEO, MXLR, USLR, and USIL), and “accessions” (as described above in *Plants and Insects*) nested within plant type. Throughout, the dependent variables were metabolite concentrations (pmol/g of fresh tissue), whether constitutive, total induced or net induced for the 12 metabolites considered in this study. *A priori* contrasts were used for paired comparisons between BTEO and MXLR (i.e., to infer domestication effects), MXLR and USLR (i.e., spread effects), and USLR and USIL (i.e., breeding effects). The significance level for contrasts was adjusted, per Sidak’s correction, to *P* ≤ 0.017 for each of the three comparisons, to maintain Type I error at or below α = 0.05 [63]. Pearson’s correlations of canonical scores with dependent variables (i.e., metabolite concentration) were used to determine the contributions of each dependent variable to the total variation in the canonical axes of MANOVA’s centroid plots; only correlations with *r* values ≥ |0.50|, and *P* ≤ 0.05 were considered significant. An additional, nested MANOVA with independent variables “plant type” and “herbivory” nested within plant type was performed on total induced (as described above) metabolite concentrations to assess whether WCR feeding affected total induced metabolite levels within each of the plant types; eta-squared (η_p_^2^) effect sizes were calculated for each of the plant types as an additional measure of how strongly seedling metabolite levels were affected by WCR feeding [64]. A critical *P* value of 0.013, per Sidak’s correction, was used for each of four *a priori* comparisons in the nested MANOVA.

Univariate analysis of variance (ANOVA) was performed to assess domestication, spread and breeding effects for all individual metabolites (constitutive, total induced, and net induced), with the independent variable “plant type” as described above for MANOVA. A nested ANOVA (as described above for MANOVA) was applied for each metabolite to compare between constitutive (i.e., free of WCR feeding) and total induced (i.e., with WCR feeding) metabolite variables within each plant type. *A priori* contrasts with critical significances of *P* ≤ 0.017 and *P* ≤ 0.013, for three and four comparisons, respectively, were used as described above. One sample *t*-tests were applied to net induced metabolite levels within plant types with the null hypothesis that levels would not differ from zero (i.e., H_0_ = 0) and alternative hypothesis that levels would be below zero (i.e., H_A_ < 0); the critical significance level was set as *P* ≤ 0.013 per Sidak’s correction [63]. All data, were transformed to meet normality assumptions prior to analyses; the Yeo IKJohnson RA [65] power transformation ln(*x* + 1) for *x* ≥ 0 or -ln(-*x* + 1) for *x* < 0 was used for analyses of net induced metabolite levels of negative values. All analyses were conducted using JMP^®^ Pro 14.0.0 software [66].

## RESULTS

### Constitutive metabolite levels

The MANOVA for levels of the constitutive metabolites among plant types showed a strong, significant multivariate effect of plant type (Wilks’ λ = 0.017, *P* < 0.001), but not of accession nested within plant type (λ = 0.008, *P* = 0.366). Variation along the *y* axis (Canonical 1) accounted for 81% of total variation, while variation along the *x* axis (Canonical 2) accounted for 17% of total variation (Figure 2A). *A priori* contrasts between pairs of plant types showed significant differences between BTEO and MXLR (F_12, 13_ = 8.950, *P* < 0.001), but not between MXLR and USLR (F_12, 13_ = 1.479, *P* = 0.205), nor between USLR and USIL (F_12, 13_ = 0.351, *P* = 0.948) (Figure 2A). Correlations of canonical scores showed that the 9-oxylipins AZA (r = 0.636, *P* < 0.001), 10-OPDA (r = −0.621, *P* < 0.001), 10-OPEA (r = −0.547, *P* < 0.001), and DA4 (r = −0.524, *P* < 0.001) accounted for the most variation along the *y*-axis, while 12-OPDA (r = −0.564, *P* < 0.001) of the 13-oxylipins branch, and AZA (r = 0.529, *P* < 0.001) accounted for most of the variation along the *x*-axis (Figure 2A).

**Figure 2.**
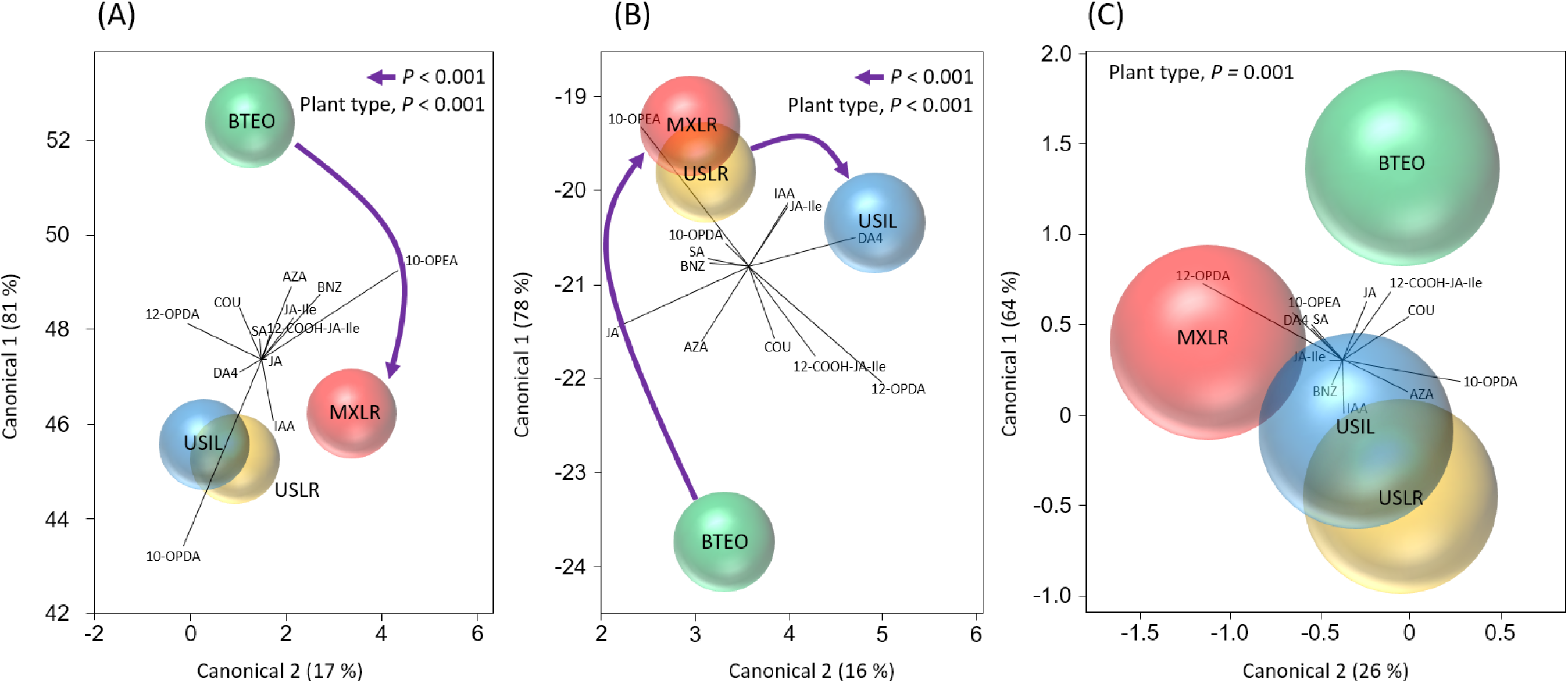
Multi-variate analysis of variance (MANOVA) centroid bi-plots for effects of plant type [Balsas teosinte (BTEO), Mexican maize landraces (MXLR), US maize landraces (USLR), and US maize inbred lines (USIL)] on levels of (**A**) constitutive, (**B**) total induced and (**C**) net induced defence-related metabolites in seedlings; corresponding *Wilks* statistics are λ = 0.017, *P* < 0.001, λ = 0.109, *P* < 0.001, and λ = 0.475, *P =* 0.001. Constitutive = seedlings not exposed to Western corn rootworm, total induced = seedlings exposed to WCR, and net induced = total induced – constitutive. In each bi-plot, spheres represent 95% confidence intervals surrounding the multivariate mean for each plant type. The independent variables included in the model were “plant type” (BTEO, MXLR, USLR, USIL), and “plant accessions” nested within plant type (not shown here). The dependent variables are shown as vectors and include the 12 metabolites considered in this study: 12-OPDA, JA, JA-Ile, 12-COOH-JA-Ile, 10-OPDA, 10-OPEA, DA4, AZA, COU, BNZ, SA, and IAA (see Figure 1 and text for full metabolite names). A solid arrow indicates a significant *a priori* contrast comparison (critical *P* ≤ 0.017, per Sidak’s correction) between plant types representing the domestication (BTEO vs. MXLR), spread (MXLR vs. USLR), and breeding (USLS vs. USIL) transitions; absence of an arrow indicates a non-significant comparison.

ANOVA performed on the constitutive levels of the 12 metabolites showed significant differences among plant types only for oxylipins, specifically for 12-OPDA and JA of the 13-oxylipins branch, and all four metabolites of the 9-oxylipins branch, (F_3, 24_ ≥ 3.173, *P* ≤ 0.042) (Table 2, Figure 3, 4). *A priori* contrasts revealed significant domestication effects (F_1, 24_ ≥ 6.665, *P* ≤ 0.016) for 12-OPDA of the 13-oxylipins branch (Figure 3A), and 10-OPDA, 10-OPEA, and DA4 of the 9-oxylipins branch (Figures 3E, F, G, 4) (Table 3). Specifically, the level of 12-OPDA decreased, while the levels of the three 9-oxylipins metabolites increased with domestication (Table 3, Figure 4). Significant spread effects (F_1, 24_ ≤ 6.634, *P* ≥ 0.017) were detected for JA and AZA of LOX (Figure 3B, H), with the level of the former increasing, and the latter decreasing (Table 3; Figure 4). Significant breeding effects were not detected for any of the 12 metabolites (Table 3, Figure 3, 4).

**Table 2.**
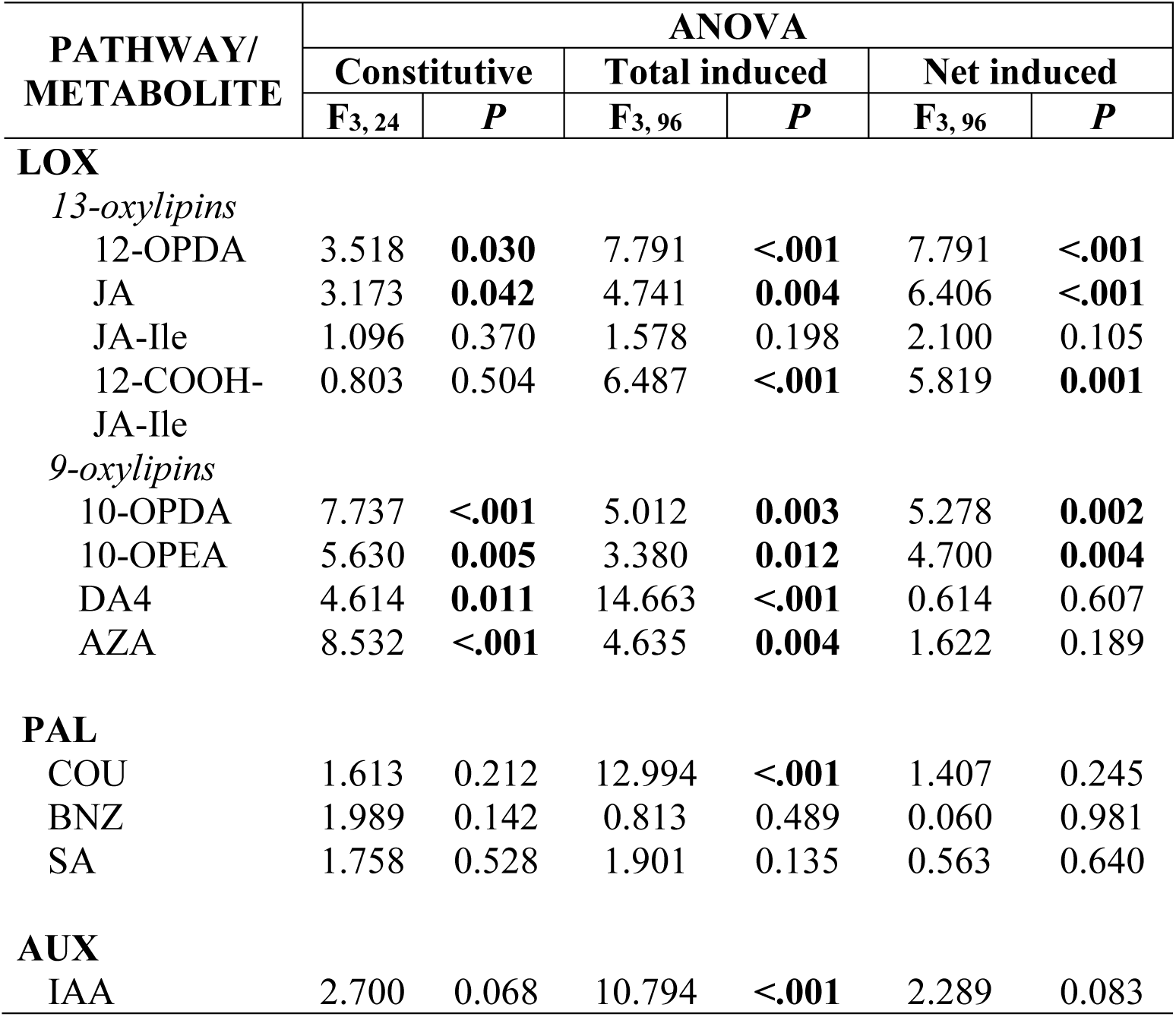
Univariate analysis of variance (ANOVA) statistics for the effect of plant type (Balsas teosinte, Mexican maize landraces, US maize landraces, US maize inbred lines) on constitutive (= without exposure to Western corn rootworm), total induced (with exposure to WCR), and net induced (= total induced level – constitutive level) levels of 12 defence-related metabolites. LOX is the lipoxygenase pathway, PAL is the phenylpropanoid pathway, and AUX is the auxin biosynthesis pathway. See Figure 1 and text for full metabolite names. Significant *P* values for ANOVA (*P* ≤ 0.05) are shown in bold type.

**Table 3.** *A priori* contrast statistics for comparisons between mean levels of defence-related metabolites (pmol/g fresh weight) in pairs of four plant types (Balsas teosinte, Mexican maize landraces, US maize landraces, US maize inbred lines) encompassing the domestication, spread, and breeding processes of maize; comparisons are shown only for metabolites shown to be affected by plant type, as shown in Table 2. Metabolite levels were measured as constitutive (= without exposure to Western corn rootworm), total induced (= with exposure to WCR), and net induced (= total induced level – constitutive level). See Figure 1 and text for full metabolite names. Significant contrasts (critical *P* ≤ 0.017, per Sidak’s correction) are indicated by bold-type.

| PATHWAY/<br>METABOLITE | CONTRASTS |  |  |  |  |  |
| --- | --- | --- | --- | --- | --- | --- |
|  | Domestication |  | Spread |  | Breeding |  |
|  | F | P<sup>1</sup> | F | P | F | P |
| **CONSTITUTIVE (*df* = 1, 24)** |  |  |  |  |  |  |
| **LOX** |  |  |  |  |  |  |
| *13-oxylipins* |  |  |  |  |  |  |
| 12-OPDA | 6.665 | **0.016**↓ | 4.839 | 0.038 | 0.576 | 0.455 |
| JA | 0.002 | 0.961 | 7.235 | **0.013**↑ | 2.558 | 0.123 |
| *9-oxylipins* |  |  |  |  |  |  |
| 10-OPDA | 17.183 | **<.001**↑ | 0.059 | 0.810 | 0.039 | 0.846 |
| 10-OPEA | 12.229 | **0.002**↑ | 0.037 | 0.848 | 0.003 | 0.955 |
| DA4 | 9.211 | **0.006**↑ | 0.003 | 0.960 | 0.009 | 0.925 |
| AZA | 2.558 | 0.123 | 6.634 | **0.017**↓ | 0.001 | 0.970 |
| **TOTAL INDUCED (*df* = 1, 96)** |  |  |  |  |  |  |
| **LOX** |  |  |  |  |  |  |
| *13-oxylipins* |  |  |  |  |  |  |
| 12-OPDA | 47.844 | **<.001**↓ | 2.133 | 0.147 | 9.579 | **0.002**↑ |
| JA | 11.914 | **<.001**↓ | 8.355 | **0.005**↑ | 1.833 | 0.179 |
| 12-COOH-JA-Ile | 11.539 | **0.001**↓ | 0.283 | 0.596 | 7.816 | **0.006**↑ |
| *9-oxylipins* |  |  |  |  |  |  |
| 10-OPDA | 9.435 | **0.003**↑ | 0.016 | 0.899 | <0.001 | 0.984 |
| 10-OPEA | 8.417 | **0.005**↑ | 0.149 | 0.700 | 0.100 | 0.756 |
| DA4 | 21.096 | **<.001**↑ | 0.687 | 0.409 | 7.156 | **0.009**↑ |
| AZA | 9.556 | **0.003**↓ | 0.006 | 0.940 | <0.001 | 0.993 |
| **PAL** |  |  |  |  |  |  |
| COU | 35.507 | **<.001**↓ | 2.006 | 0.160 | 0.433 | 0.512 |
| **AUX** |  |  |  |  |  |  |
| IAA | 6.621 | **0.012**↑ | 2.179 | 0.143 | 1.908 | 0.170 |
| **NET INDUCED (*df* = 1, 96)** |  |  |  |  |  |  |
| **LOX** |  |  |  |  |  |  |
| *13-oxylipins* |  |  |  |  |  |  |
| 12-OPDA | 0.119 | 0.730 | 15.974 | **<.001**↓ | 6.535 | **0.012**↑ |
| JA | 5.310 | 0.023 | 3.396 | 0.068 | 0.733 | 0.394 |
| 12-COOH-JA-Ile | 11.733 | **<.001**↓ | 0.079 | 0.779 | 0.636 | 0.427 |
| *9-oxylipins* |  |  |  |  |  |  |
| 10-OPDA | 13.703 | **<.001**↓ | 0.385 | 0.536 | 0.298 | 0.586 |
| 10-OPEA | 9.883 | **0.002**↓ | 0.035 | 0.852 | 0.843 | 0.361 |

<sup>1</sup>Red, down-pointing arrows (↓) indicate a decrease, and green, up-pointing arrows (↑) an increase for a particular metabolite.

**Figure 3.**
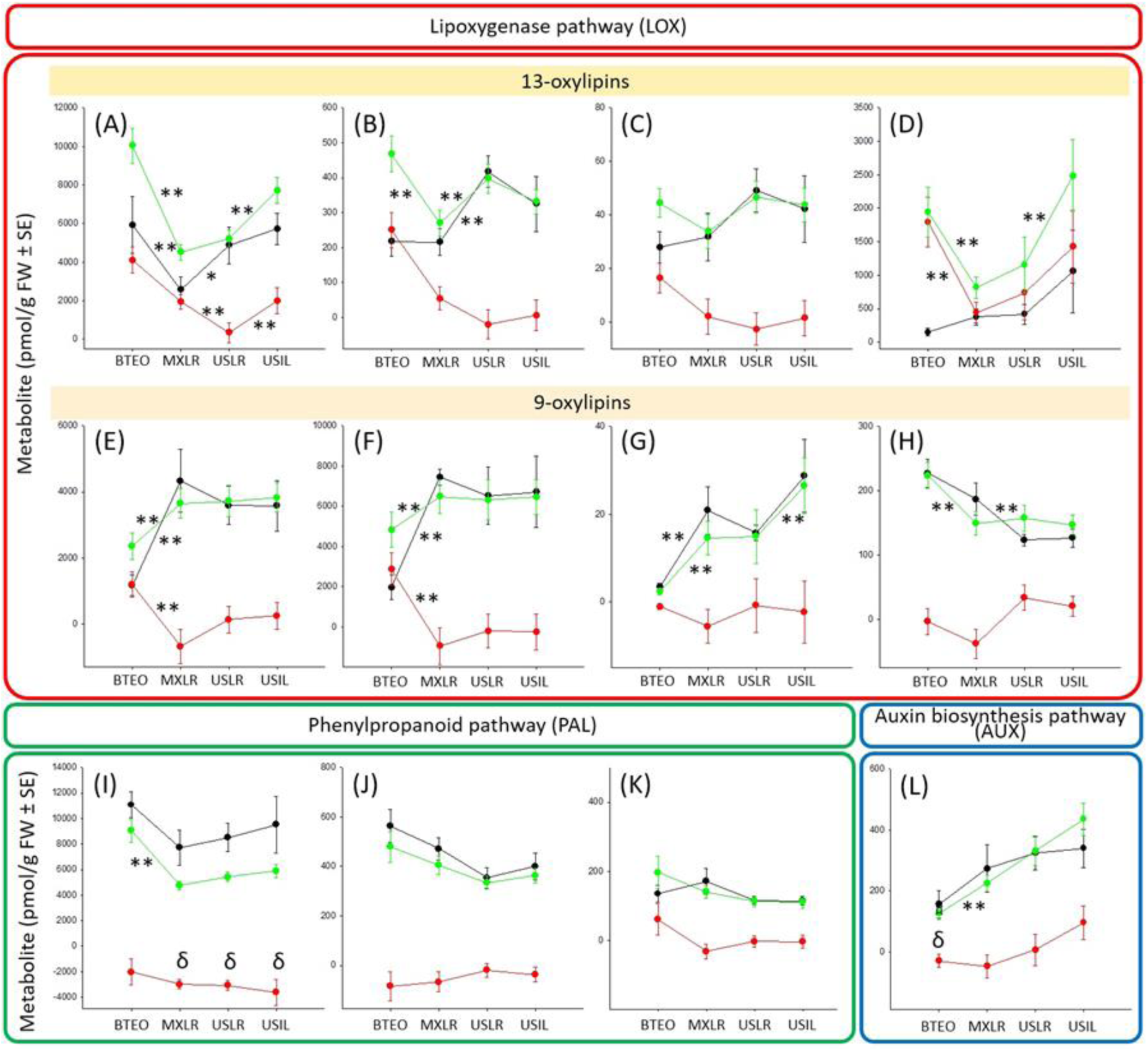
Mean levels of 12 defence-related metabolites (pmol/g fresh weight) in four plant types ordered from most ancestral to most derived: Balsas teosinte (BTEO), Mexican maize landraces (MXLR), US maize landraces (USLR), and US maize inbred lines (USIL). The metabolites are: **(A)** 12-OPDA, **(B)** JA, **(C)** JA-Ile, **(D)** 12-COOH-JA-Ile, **(E)** 10-OPDA, **(F)** 10-OPEA, **(G)** DA4, **(H)** AZA, **(I)** COU, **(J)** BNZ, **(K)** SA, and **(L)** IAA (see Figure 1 and text for full metabolite names). Black lines represent constitutive metabolite levels (without exposure to Western corn rootworm), green lines represent total induced levels (with exposure to WCR), and red lines represent net induced levels (= total induced level – constitutive level). Asterisks indicate significant *a priori* contrasts between plant types representing domestication (BTEO vs. MXLR), spread (MXLR vs. USLR), and breeding (USLR vs. USIL), * *P* ≤ 0.05, ** *P* ≤ 0.001; corresponding ANOVA statistics are shown in Table 2, and contrast statistics in Table 3. Plots are grouped by metabolic pathway, as shown by inset labels and coloured outlines. For net induced levels δ, indicates mean value significantly below zero (*t ≥* 2.840, 26 d.f., *P* ≤ 0.004; critical *P* ≤ 0.013, per Sidak’s correction).

**Figure 4.**
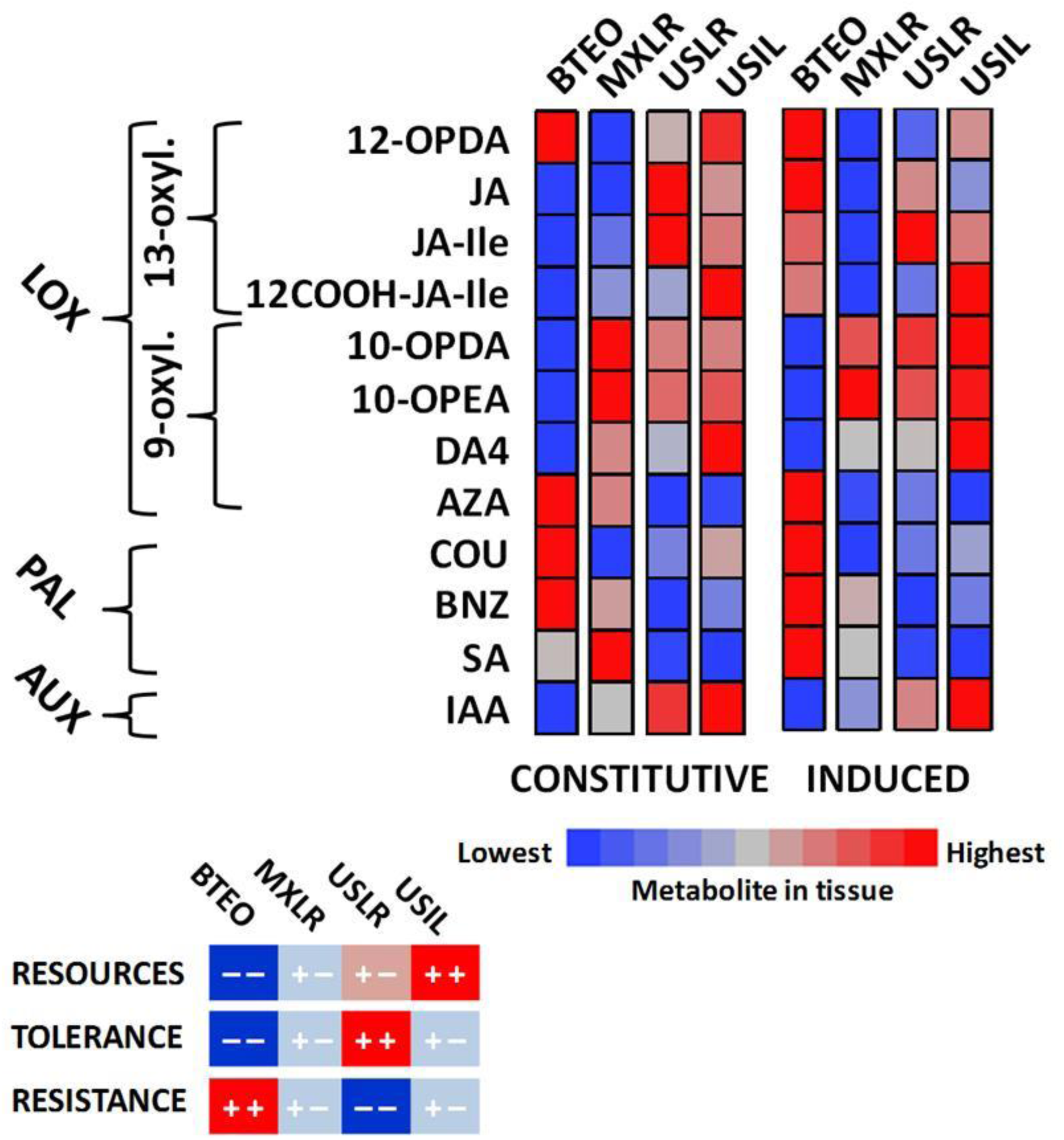
Heat map for constitutive and total induced levels of 12 metabolites in roots of seedlings of four plant types [Balsas teosinte (BTEO), Mexican maize landraces (MXLR), US maize landraces (USLR), US maize inbred lines (USIL)]; respectively, constitutive and total induced metabolite levels correspond to seedlings not exposed or exposed to feeding by Western corn rootworm. For each metabolite and across plant types (i.e. horizontally), blue coloration indicates the lowest metabolite level and red indicates the highest level (see heat legend at bottom). Overall effects of domestication, spread, and breeding on metabolite levels are suggested by differences between plant types within each group of columns (Constitutive, Total induced); overall effects of feeding by Western corn rootworm on metabolite levels for each plant type are suggested by differences between corresponding bars between groups of columns. The relative levels of resource richness, and resistance and tolerance to Western corn rootworm for each plant type are shown inset at the bottom of the figure (from Fontes-Puebla and Bernal, 2020).

Overall, these results suggested a strong effect of domestication on constitutive levels of 9-oxylipins and the 13-oxylipin 12-OPDA, strong to modest effects of spread on LOX metabolites, and minimal to no effects of breeding on any metabolites (Table 3, Figure 3, 4). Specifically, MANOVA revealed only a domestication effect (Figure 2A), and ANOVA showed that constitutive levels of three 9-oxylipins, 10-OPDA, 10-OPEA, and DA4, were associated with domestication (Figure 3E-G), and the 13-oxylipin 12-OPDA decreased with domestication (Figure 3A) (Table 3). Spread affected only the levels of the 13-oxylipins 12-OPDA and JA (Figure 3A, B), and the 9-oxylipin AZA (Figure 3H), which increased and decreased, respectively (Table 3); breeding did not affect the levels of any of the 12 metabolites (Table 3, Figure 3).

### Total induced metabolite levels

MANOVA on levels of total induced metabolites showed significant effects of plant type (λ = 0.109, *P* < 0.001) and accession nested within plant type (λ = 0.146, *P* < 0.001).Variation along the *y* axis (Canonical 1) accounted for 78% of total variation, while variation along the *x* axis (Canonical 2) accounted for 16% of total variation (Figure 2B). *A priori* contrasts between plant types indicated differences between BTEO and MXLR (F_12, 85_ = 19.794, *P* < 0.001), and between USLR and USIL (F_12, 85_ = 4.092, *P* < 0.001), but not between MXLR and USLR (F_12, 85_ = 1.947, *P* = 0.040) (Figure 2B). Correlation of canonical scores showed that the 13-oxylipin 12-OPDA (r^2^ = −0.608, *P* < 0.001), the 9-oxylipin DA4 (r^2^ = 0.520, *P* <0.001), and the phenylpropanoid COU (r^2^ = −0.593, *P* <0.001) contributed significantly to variation along the *y* axis, while no metabolite contributed significantly (i.e., *r* ≥ |0.50|, *P* ≤ 0.05) to variation along the *x* axis (Figure 2B).

ANOVA performed on the total induced levels of the 12 metabolites showed significant differences among plant types for all oxylipins, except JA-Ile, as well as for COU, and IAA (F_3, 96_ ≥ 3.380, *P* ≤ 0.012) (Table 2, Figure 3, 4). *A priori* contrasts between pairs of plant types revealed significant domestication effects (F_1, 96_ ≥ 8.417, *P* ≤ 0.005) for all LOX metabolites, except JA-Ile, with 13-oxylipins consistently decreasing (Figure 3A, B, D, 4), 9-oxylipins consistently increasing (Figure 3E-G, 4), except AZA, which decreased (Figure 3H, 4) (Table 3); also, COU decreased (Figure 3I, 4), and IAA increased (Figure 3L, 4) with domestication (F_1, 96_ ≥ 6.621, *P* ≤ 0.012) (Table 3). Additionally, *a priori* contrasts revealed that JA increased with spread (Figure 3B, 4), while 12-OPDA, 12-COOH-JA-Ile, and DA4 increased with breeding (Figure 3A, D, G, 4) (F_1, 96_ ≥ 7.156, *P* ≤ 0.009) (Table 3).

Overall, these results suggested that total induced levels of oxylipins, in particular, were strongly affected by domestication, minimally affected by spread, and modestly affected by breeding (Table 3, Figure 4). Specifically, MANOVA revealed significant domestication and breeding effects (Figure 2B). Concomitantly, ANOVA showed that domestication decreased the total induced levels of the 13-oxylipins 12-OPDA, JA, and 12-COOH-JA-Ile (Figure 3A, B, D), and increased the levels of all 9-oxylipins, except of AZA, which increased (Figure 3H); the level of the PAL metabolite COU decreased, and of IAA increased(Figure 3I, L) (Table 3, Figure 4). Spread affected only the level of JA, which increased, while breeding affected the levels of 12-OPDA and 12-COOH-JA-Ile, and DA4, which increased (Figures 3B, E-L) (Table 3, Figure 4).

### Net induced metabolite levels

MANOVA on the levels of net induced metabolites (i.e., constitutive minus total induced metabolite levels) showed a strong, significant multivariate effect for plant type (Wilks’ λ = 0.475, *P* = 0.001), and for accession nested within plant type (λ = 0.141, *P* < 0.001); variation along the *y* axis (Canonical 1) accounted for 64% of total variation, while variation along the *x* axis (Canonical 2) accounted for 26% of total variation (Figure 2C). *A priori* contrasts between plant type multivariate means showed no significant differences between BTEO and MXLR (F_12, 85_ = 2.090, *P* = 0.026), between MXLR and USLR (F_12, 85_ = 1.942, *P* = 0.040), nor between USLR and USIL (F_12, 85_ = 1.061, *P* = 0.403) (Figure 2C). Correlation analyses showed that the 13-oxylipins 12-OPDA (*r*^*2*^ = 0.614, *P* < 0.001), JA (*r*^*2*^ = 0.683, *P* < 0.001), and 12-COOH-JA-Ile (*r*^*2*^ = 0.604, *P* < 0.001), and the 9-oxylipin 10-OPEA (*r*^*2*^ = 0.524, *P* < 0.001), contributed significantly to the separation among plant type multivariate means along the *y* axis (Figure 2C).

ANOVA performed on the net induced levels of the 12 metabolites showed significant differences among plant types, though only for the 13-oxylipins 12-OPDA, JA, and 12-COOH-JA-Ile, and the 9-oxylipins 10-OPDA and 10-OPEA (F_1, 96_ ≥ 4.700, *P* ≤ 0.004) (Table 2, Figure 3). *A priori* contrasts showed significant (F_1, 96_ ≥ 9.883, *P* ≤ 0.002) reductions with domestication for JA, 12-COOH-JA-Ile, 10-OPDA, and 10-OPEA (Figure 3, B, D, E, F, Table 3, Figure 4). Significant spread and breeding effects (F_1, 96_ ≥ 6.535, *P* ≤ 0.012) were shown only for 12-OPDA, which decreased and increased, respectively (Figure 3A) (Table 3, Figure 4). One sample *t*-tests showed that mean levels of COU were significantly below zero for all plant types, except BTEO (Figure 3I), and mean level of IAA was below zero for BTEO (Figure 3L) (*t* ≥ 2.840, *P* ≤ 0.004), suggesting that these metabolites were suppressed by herbivory.

Overall, these results suggested modest effects of domestication, evident as effects on net induced levels of oxylipins, and minimal effects of spread and breeding, evident as effects on the levels of 12-OPDA. Specifically, while MANOVA did not reveal significant effects of domestication, spread or breeding (Figure 2C), ANOVA showed that domestication was associated with decreased net induced levels of the oxylipins 12-COOH-JA-Ile, 10-OPDA, and 10-OPEA (Figures 3B, D, E, F), while spread decreased and breeding increased the level of 12-OPDA (Figure 3A) (Table 3, Figure 4). Additionally, net induced levels of COU were significantly below zero in all plant types, except BTEO (Figure 3I), while the net induced level of IAA was below zero only in BTEO (Figure 3L).

### Effects of WCR herbivory on individual plant types

MANOVA on levels of total induced metabolites showed significant effects of plant type (λ = 0.223, P < 0.0001) and WCR nested within plant type (λ = 0.482, P < 0.0001). Variation along the *y* axis (Canonical 1) accounted for 73% of total variation, while variation along the *x* axis (Canonical 2) accounted for 16% of total variation (Figure 5A). *A priori* contrasts within plant types, i.e., between WCR-infested (= total induced) and non-infested (= constitutive) levels, showed significant differences for BTEO (F_12, 129_ = 3.905, P < 0.0001) and MXLR (F_12, 129_ = 2.723, P = 0.003), but not for USLR (F_12, 129_ = 1.230, P = 0.270), and USIL (F_12, 129_ = 1.327, P = 0.212) (Figure 5A). The effect of WCR herbivory was strongest in BTEO (η_p_^2^ = 0.274), followed by MXLR (η_p_^2^ = 0.209), USIL (η_p_^2^ = 0.114), and USLR (η_p_^2^ = 0.106). This result is evident in Figure 4 as clearly contrasting heat bars between WCR(−) and WCR(+) treatments for BTEO, modestly contrasting for MXLR, and minimally contrasting for USLR and USIL. Correlation of canonical scores showed that the 13-oxylipins 12-OPDA (r^2^ = –0.695, *P* < 0.0001) and 12-COOH-JA-Ile (r^2^ = –0.608, *P* <0.0001) contributed significantly to variation along the *y* axis; no metabolite was found to contribute significantly (i.e., *r* ≥ |0.50|, *P* ≤ 0.05) to variation along the *x* axis (Figure 5A).

**Figure 5.**
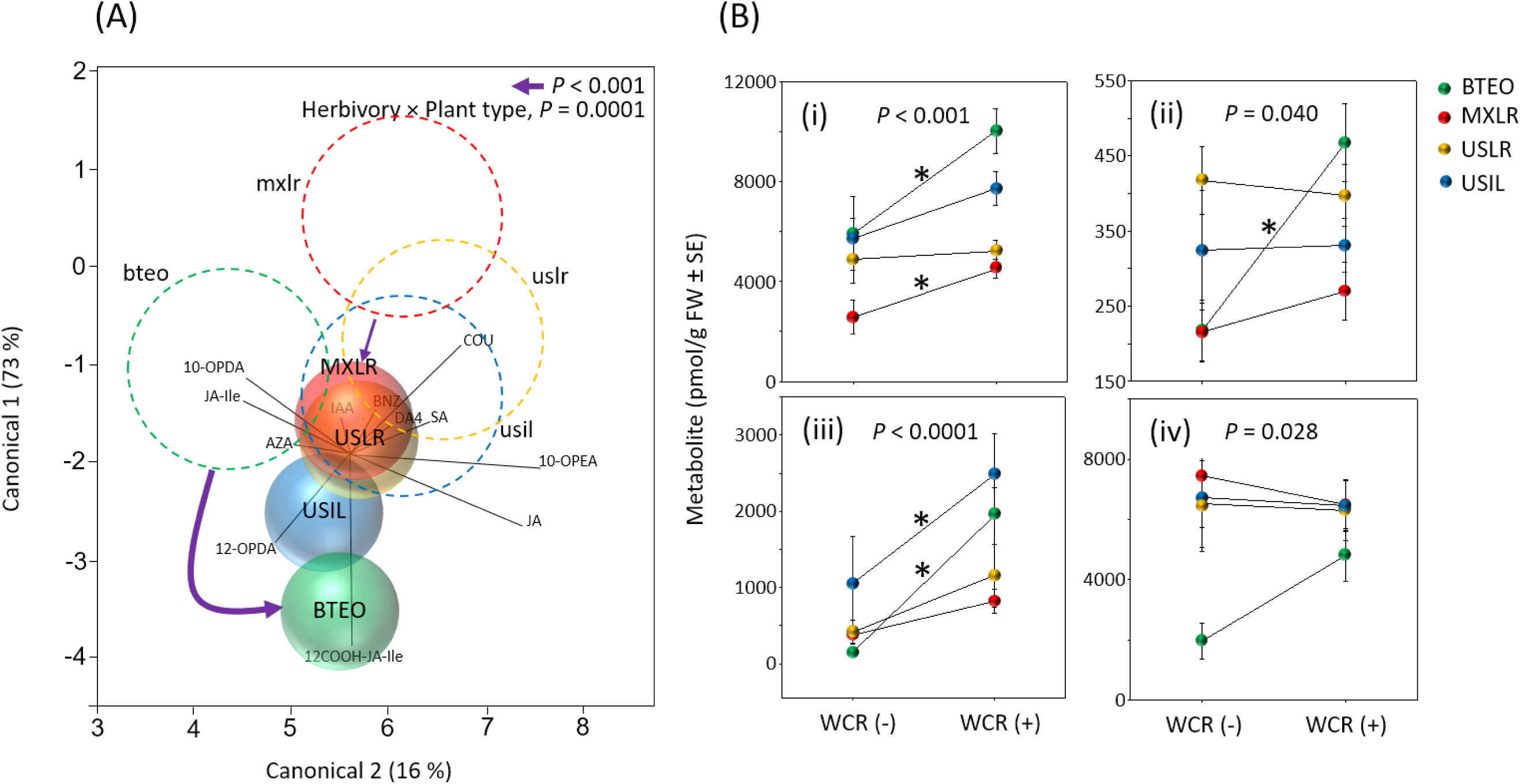
**(A)** MANOVA centroid bi-plot for defence-related metabolites in seedlings of four plant types [Balsas teosinte (BTEO), Mexican maize landraces (MXLR), US maize landraces (USLR), US maize inbred lines (USIL)] as affected by feeding by Western corn rootworm (WCR) (Wilks λ = 0.482, *P* < 0.0001). Respectively, solid spheres and dashed circles represent 95% confidence intervals around multivariate means for WCR-infested (labelled with upper-case text) and non-infested (lower-case text) seedlings. The model included the nested term “herbivory [plant type]” (herbivory = WCR-infested or non-infested; plant type = BTEO, MXLR, USLR, USIL). The dependent variables are shown as vectors, and included: 12-OPDA, JA, JA-Ile, 12-COOH-JA-Ile, 10-OPEA, 10-OPDA, DA4, AZA, COU, BNZ, SA, and IAA (see Figure 1 and text for full metabolite names). The solid arrows indicate significant *a priori* contrasts (critical *P* ≤ 0.013, per Sidak’s correction) between WCR-infested and non-infested Balsas teosinte seedlings (arrow width is proportional to the significance level); the absence of an arrow between plant type pairs indicates a non-significant contrast. **(B)** Nested-effect plots for significant ANOVAs (F_3, 140_ ≥ 2.775, *P* ≤ 0.05) depicting effects of feeding by WCR [without feeding = WCR(-); with feeding = WCR(+)] on mean metabolite levels (pmol/g of fresh weight ± SE) in seedlings of the four plant types: **(i)** 12-OPDA (F_4,136_ = 5.821, *P* < 0.001), **(ii)** JA (F_4,136_ = 2.589, *P* = 0.040), **(iii)** 12-COOH-JA-Ile (F_4,136_ = 6.629, *P* < 0.0001), **(iv)** COU (F_4,136_ = 2.819, *P* = 0.028). Asterisks indicate significant contrasts (critical *P* ≤ 0.013, per Sidak’s correction).

The nested term in the ANOVA (herbivory within plant type) of total induced levels of each of the 12 metabolites showed significant effects of herbivory on levels of the 13-oxylipins 12-OPDA, JA, and 12-COOH-JA-Ile (F_4, 136_ ≥ 2.589, P ≤ 0.040), and the phenylpropanoid COU (F_4, 136_ = P = 0.028), but not on the remaining metabolites (F_4, 136_ ≤ 1.861, P ≥ 0.121) (Table 4). In particular, increases with herbivory were evident in 12-OPDA, JA, 12-COOH-JA-Ile in BTEO (P ≤ 0.002), 12-OPDA in MXLR (P = 0.001), and 12-COOH-JA-Ile in USIL (P = 0.013) (Figure 5B); while COU increased minimally in each plant type, the increases were not significant (F_1, 135_ ≤ 4.350, P = 0.039).

**Table 4.** Nested analysis of variance (ANOVA) statistics for effects of plant type (Balsas teosinte, Mexican maize landraces, US maize landraces, US maize inbred lines) on levels of defence-related metabolites in seedlings exposed or not exposed to Western corn rootworm. LOX is the lipoxygenase pathway, PAL is the phenylpropanoid pathway, and AUX is the auxin biosynthesis pathway. See Figure 1 and text for full metabolite names. Significant effects of infestation by Western corn rootworm (*P* ≤ 0.05) are shown in bold.

| PATHWAY/<br>METABOLITE | NESTED ANOVA |  |
| --- | --- | --- |
|  | F<sub>4, 136</sub> | P |
| **LOX** |  |  |
| *13-oxylipins* |  |  |
| 12-OPDA | 5.821 | **<.001** |
| JA | 2.589 | **0.040** |
| JA-Ile | 0.627 | 0.644 |
| 12-COOH-JA-Ile | 6.629 | **<.0001** |
| *9-oxylipins* |  |  |
| 10-OPDA | 0.861 | 0.489 |
| 10-OPEA | 1.821 | 0.128 |
| DA4 | 1.861 | 0.121 |
| AZA | 0.753 | 0.558 |
| **PAL** |  |  |
| COU | 2.819 | **0.028** |
| BNZ | 0.170 | 0.953 |
| SA | 1.010 | 0.405 |
| **AUX** |  |  |
| IAA | 0.538 | 0.708 |

Altogether, these results suggested that the overall defence response to WCR feeding (= total induced metabolite levels) was strongest in BTEO, modest in MXLR, and minimal in USLR and USIL. Specifically, MANOVA revealed significant defence responses to WCR for BTEO and MXLR (Figure 5A), and ANOVA showed that for BTEO, most of the 13-oxylipins significantly increased with WCR feeding (Figure5B), while single metabolites increased in MXLR (12-OPDA) and USIL (12-COOH-JA-Ile). Put together, our results suggested that BTEO responds to WCR injury by increasing its production of 13-oxylipins, while maintaining 9-oxylipins and IAA at low levels (Figure 4). In contrast, the remaining plant types showed weak to minimal responses to WCR feeding (Figure 4).

## DISCUSSION

This study addressed whether the production of selected phytohormones and metabolites relevant to resistance and tolerance to Western corn rootworm in maize were mediated by the crop’s domestication, northward spread, and modern breeding, processes spanning thousands of maize and WCR generations, and divergent environments across ∼20 ° latitude and >3000 m elevation. We hypothesized that we would find: (i) a shift from reliance on induced to constitutive herbivore defence with domestication and cultivation in increasingly rich environments; (ii) a decrease in levels of defence-related oxylipins, and; (iii) an increase in the growth-promoting phytohormone, IAA [37, 38, 40, 54].

Consistently with our hypotheses, we found that metabolite levels were strongly affected by WCR feeding in Balsas teosintes, modestly in Mexican landraces, and minimally in US landraces and inbred lines (see Figure 4, 5), indicative of a shift from induced to constitutive herbivore defence with domestication and cultivation in increasingly rich environments. Also, we found changes in levels of oxylipin metabolites associated with domestication, spread, and breeding, though their direction differed between 13- and 9-oxylipin metabolites: Constitutive levels of 9-oxylipins increased with domestication, but minimally changed with spread and breeding, while those of 13-oxylpins seemed to decrease modestly with domestication and recover with spread and breeding (see Figure 3, 4). Finally, levels of the growth-promoting phytohormone, IAA, increased with domestication, followed by an increasing trend with spread and breeding, and cultivation in increasingly rich environments (Figure 3, 4).

### With domestication, maize resistance to root herbivory shifted from reliance on induced to reliance on constitutive defence responses

Balsas teosinte, but not maizes, seemed to rely on activation of the 13-LOX pathway for root defence against WCR. Feeding by WCR larvae consistently increased most 13-oxylipins in BTEO (except JA-Ile), while causing increases in the levels of only single 13-oxylipins in MXLR (12-OPDA) and in USIL (12-COOH-JA-Ile) (see Figure 4, 5). In contrast, constitutive (and total induced) levels of 9-oxylipins (except AZA), but not 13-oxylipins, were consistently higher in MXLR compared to BTEO (see Figure 3), which suggests that the 9-LOX pathway was under positive selection during domestication. This observation is in line with recent evidence showing the 9-LOXs ZmLOX3 and its orthologue to be under parallel selection in both maize and sorghum during their domestication [67]. In addition to their roles as signals, 10-OPEA and derivatives also possess strong insecticidal activity [22], and although the net induced levels of these 9-oxylipins were consistently near nil in the maizes, the corresponding induced levels of 10-OPDA and 10-OPEA were significantly higher in BTEO compared to MXLR (see Figure 3). While previous studies showed that 10-OPEA increased in maize roots 7 days after WCR feeding, which was correlated with increases in LOX3 and other 9-LOXs [68], our data suggests that, relative to the maizes, BTEO can more readily activate this sub-branch of the 9-LOX pathway in response to WCR feeding. Coincident with this study’s results, our previous study showed that BTEO was more resistant but less tolerant to WCR than MXLR and maizes generally [51]. Put together, this and our prior study’s results suggest that the stronger resistance and weaker tolerance to WCR in BTEO compared to MXLR is likely due to elevated levels of induced jasmonates and low levels of IAA in BTEO, while defence in MXLR, and maize generally, is due to high constitutive levels of 9-oxylipins and IAA.

Aside from the oxylipins, changes from domestication were observed with the phenylpropanoid COU and the auxin IAA, which were lower and higher, respectively, in MXLR compared to BTEO. Intriguingly, net induced levels of COU were generally below zero in the maizes (i.e. total induced levels were lower than constitutive levels), which raises the possibility that COU is used as substrate for non-salicylate phenylpropanoid [69], especially given that SA levels are not dramatically changed in maize roots following WCR feeding (see Figure 3K, 4; [33]). Similar to LOX3, the phenylpropanoid biosynthetic enzymes, PAL4 and PAL6, also appear to be under selection pressure during the domestication of maize [67].

Given the lower costs of induced compared to constitutive defences [36, 70], our results are consistent with the hypothesis that plants in high-resource environments should rely predominantly on constitutive defences because they are not physiologically constrained, while those adapted to low-resource environments should rely on induced defences. A gradient of resource availability is shown in Figure 4, suggesting that environmental resources are poorest for Balsas teosinte and richest for US inbred lines. Teosintes (*Zea* spp., except maize) generally grow in marginal environments that are unsuitable for agriculture and livestock [71, 72]. Indeed, the three Balsas teosinte accessions included in this study were collected from thin, sandy soils on mountain slopes (El Rodeo, Talpitita) or along a roadside (Ejutla) (JSB, unpubl.). For their part, Mexican landraces are grown in poorer soils compared to improved commercial varieties. Prior studies showed that landrace maizes are planted on ∼80% of the area devoted to maize in Mexico, encompassing mostly hillsides with poor soils, while improved commercial varieties are planted on the remaining ∼20% of the area, and that farmers will plant commercial varieties on good soils and landraces on poorer soils, when they have a choice [73-77]. Finally, while US landraces (no longer planted commercially) and inbred lines were developed for the same environments, the availability of major soil nutrients, e.g., nitrogen, differed greatly between the early and late portions of the 20^th^ century, associated with intensification of maize production reliant on commercial (hybrid) varieties and other inputs (e.g., fertilizers and insecticides to support high crop productivity and offset insect injury) that began in the middle of that century [50, 51, 78, 79]. For example, prior to the 1930s, few maize growers employed fertilizers because they were uneconomical or unavailable, while the use of synthetic nitrogen fertilizers nearly tripled between 1964 and 1985 [78-80].

### Domestication engaged activation of 9-oxylipins and diminished role of 13-oxylipins during maize defence against WCR

Our results suggested that while levels of 13-oxylipins (except JA-Ile) decreased with domestication, they tended to recover with breeding (12-OPDA, 12-COOH-JA-Ile) or spread (JA) (see Figure 3). In contrast, levels of 9-oxylipins (except AZA) appeared to increase to a plateau with domestication, with no further changes (10-OPDA, 10-OPEA), or with an increase with breeding (DA4) (see Figure 3). 12-OPDA showed the highest constitutive and total induced levels among the 13-oxylipins, and is a JA precursor that has a signalling role distinct from the canonical phytohormone JA-Ile [81]. Recent studies supported such a unique role for 12-OPDA in maize by showing that it, but not JA, is a major defence signal against sap-sucking aphids [82], and a long-distance signal in symbiont-induced systemic resistance against pathogens [23]. Total induced levels of 12-OPDA were highest in BTEO, intermediate in USIL, and lowest in MXLR and USLR.

Jasmonate catabolism breaks down JA-Ile, the form of JA that is bioactive and ultimately responsible for plant defence responses, to produce 12-COOH-JA-Ile and reduce metabolic expenditures, though whether this metabolite has its own activity or is merely a catabolite of biologically active JA is unknown [1, 83-87]. Largely similar to the case of 12-OPDA, total induced levels of 12-COOH-JA-Ile were highest in BTEO and USIL, and lowest in MXLR and in USLR. Interestingly, BTEO showed high resistance to WCR, MXLR and USIL displayed intermediate resistance, and USLR had low resistance in our previous study [51], suggesting a correlation between WCR resistance and levels of 12-OPDA and 12-COOH-JA-Ile. Other studies showed that 12-OPDA enhances callose accumulation and *Maize Insect Resistance1-Cysteine Protease* in maize upon attack by corn leaf aphid (*Rhopalosiphum maidis* Fitch), and that it acts as a cell protector by contributing to maintaining homeostasis in plants during physiological stress [88]. Similarly, *Arabidopsis* and rice catabolize derivatives of 12-OPDA, such as JA and JA-Ile, to 12-OH-JA and 12-COOH-JA-Ile, respectively, as measures to reduce metabolic costs [84, 89]. Thus, the high levels of total induced 12-OPDA in BTEO may help maintain cell homeostasis upon physiological stress due to WCR feeding, and high levels of 12-COOH-JA-Ile in BTEO and USIL may indicate an extremely rapid turnover of JA-Ile to maintain growth and reduce metabolic expenditures [51, 84]. Balsas teosinte is typically relegated to marginal environments, so may be adapted to grow under physiological stress, while maize landraces and inbred lines are grown in relatively rich agricultural contexts, in which physiological stress is minimized, as discussed above. Thus, the reduction of JA-Ile levels with domestication, and increases in 12-OPDA and 12-COOH-JA-Ile levels with breeding may be consequences of artificial selection favouring growth in a context of lower physiological stress under intensive agriculture.

Constitutive and total induced levels of AZA seemed to decrease with domestication and spread. This may suggest that maize may be allocating resources towards the synthesis of other 9-oxylipin branch derivatives, instead of AZA, which in turn crosstalk with the 13-oxylipins pathway [21, 90]. Derivatives of 9-LOX have been documented to mediate induced systemic resistance, and influence root growth, among other developmental traits, as well as mediate defensive responses by acting directly as phytoalexins against pathogens [60, 91]. AZA acts as a signal associated with SAR (systemic acquired resistance), particularly an inducer of SA accumulation upon pathogen attack [92, 93]. For example, one study found that AZA plays an important role in maize response to infection by sugarcane mosaic virus [94]. However, another study proposed that AZA functions as a signal in multiple systemic immunity programs, not exclusively as an inducer of SAR, and may cause morphological changes in roots by inhibiting primary root growth and increasing lateral root density [60]. Interestingly, Balsas teosinte roots are denser than those of modern maize [95, 96]. Thus, high constitutive and total induced levels of this compound may be related to either priming against pathogens, or functions related to root growth and herbivory tolerance through growth compensation.

Domestication enhanced the synthesis of the 9-oxylipns 10-OPEA, 10-OPDA, and DA4, and the latter was enhanced further with breeding, as noted above. These 9-oxylipins derivatives were named death acids (DA) because they were associated with necrotic tissue in pathogen-infected maize, suggesting that they mediate cell death upon pathogen attack [22]. While the function of the majority of 9-oxylipins is poorly understood, some 9-oxylipins regulate JA biosynthesis, as well as cell death upon pathogen attack [91, 97]. As a result, high levels of 10-OPEA may impair insect growth, e.g., of *Helicoverpa zea* [22].

Whether DAs act as defence signals or growth regulators, increased constitutive levels of 10-OPDA, 10-OPEA, and DA4 suggest that maize allocates resources after root herbivory towards the 9-oxylipins branch, rather than the 13-oxylipins branch. Plausibly, selection for changes in plant architecture (i.e. branching in Balsas teosinte but not maize) and enhanced growth (i.e. higher productivity in maize compared to Balsas teosinte) drove this change, which may have repercussions for herbivory and disease defence. Our results showed that the constitutive and total induced levels of DA4 were highest in USIL and lowest in BTEO, while our previous study showed weaker resistance in USILs compared to BTEOs [51]. This suggests that DA4 alone may not be relevant to resistance against WCR, and that any insecticidal properties of death acids occur through the other chemical species [22, 51].

### Levels of the growth-promoting phytohormone IAA increased with maize domestication, spread, and breeding

Our results showed that levels of the growth-related hormone IAA increased with domestication, and suggested an overall increasing trend in IAA levels from BTEO to USIL (Figure 3, 4). Intriguingly, net induced levels of IAA were negative in BTEO (i.e. total induced level < constitutive level), while they were nil and greater in all three maizes, which raises the possibility that in BTEO, IAA may be suppressed by the increase in JA in response to WCR feeding [98, 99]. Wild plants, such as Balsas teosinte, tend to be more resistant to herbivory than domesticated plants, as domestication favoured fast-growth and high productivity over defence [41, 43, 47, 48, 51]. Additionally, geographical spread, and exposure to new biotic and abiotic stressors, may reshape plant defence responses [51-53, 100, 101]. Thus, increased resource availability and novel herbivory pressure following domestication and spread may have selected for fast-growth in maize, and lead to tolerance as a defence strategy [37, 51, 52]. More recently, systematic breeding in the context of agricultural intensification and increased WCR pressure [50, 51, 102, 103] seems to have selected for resistance in maize, without measurably affecting productivity [51, 80, 104]. Our previous study showed a trade-off between WCR resistance and tolerance, and particularly that US inbred lines gained modest resistance to WCR, while losing tolerance, compared to US landraces [51]. In this study, total induced levels of IAA increased linearly from BTEO to USIL, which suggests that increased levels of IAA promote tolerance through root compensation, though tolerance trades off with resistance, as shown previously [51].

## CONCLUSION

The processes studied here, domestication, spread, and breeding, especially affected the lipoxygenase and auxin biosynthesis pathways, while a single metabolite, COU, of the phenylpropanoid pathway was affected. We found that levels of COU decreased with domestication, and remained unchanged with spread and breeding. Interestingly, however, the net induced levels of COU were consistently negative (i.e. constitutive level < total induced level) in the maizes, but not BTEO, which may indicate its use as a substrate for other phenylpropanoid metabolites to strengthen cell walls or act as feeding deterrents during WCR herbivory. These findings may merit further research.

Overall, our results were broadly consistent with our hypotheses concerning the evolution of biochemical defence strategies in maize, as mediated by domestication, spread, and breeding. These processes especially affected the lipoxygenase and auxin biosynthesis pathways, and minimally affected the phenylpropanoid pathway. The first of those pathways is relevant to insect and pathogen defence, the second to plant growth, and the third to pathogen and insect defence [6, 10, 12, 20, 60, 61, 105]. Changes in levels of oxylipins and IAA seemed to underlie our broad findings. Namely, with domestication, spread and breeding, and in the context of cultivation in increasingly rich environments, we found: i) a shift from reliance on induced to constitutive herbivore defence, (ii) a decrease in levels of 13-oxylipins and increase in levels of 9-oxylipins, putative insect defence metabolites, and (iii) an increase in levels of IAA, a growth phytohormone. Finally, this and a prior study’s results put together indicate that defence evolution in crops is mediated not only by processes spanning thousands of plant and insect generations, such as crop domestication and spread, but also by processes acting over tens of generations, such as modern breeding and agricultural intensification.

## ACKNOWLEDGEMENTS

We are grateful to Chad Nielson (USDA ARS, North Central Agricultural Research Laboratory, Brookings, SD, United States) for providing Western corn rootworm eggs, Mark Millard (USDA NPGS, Ames, IA, United States) for providing Mo17, and Mexican and US maize landrace seed, and R. F. Medina and K. Zhu-Salzman (both Texas A&M University, College Station) for helping to improve an earlier version of the manuscript. This study was supported in part by funding from: CONACyT and INIFAP (both Mexico) to AF-P (CONACyT scholarship #382690); TAMU-CONACyT [Characterization of Resistance to Root-And Foliage-Feeding Insects in Maize Breeding Lines, Landraces and Wild Ancestors, Project 2014-024(S)] and USDA Hatch (TEX07234) to JSB, and; USDA-NIFA [Systemic oxylipin signals (SOS) for herbivory-induced defense, Project USDA-NIFA (2017-67013-26524)] to MVK.

## REFERENCES

1 Wasternack, C., Kombrink, E. 2010 Jasmonate: Structural Requirements for Lipid-Derived Signals Active in Plant Stress Responses and Development. ACS Chemical Biology. 5, 63–77.

2 Erb, M., Kollner, T. G., Degenhardt, J., Zwahlen, C., Hibbard, B. E., Turlings, T. C. 2011 The role of abscisic acid and water stress in root herbivore-induced leaf resistance. New Phytol. 189, 308-320. (10.1111/j.1469-8137.2010.03450.x)

3 Maes, L., Goossens, A. 2010 Hormone-mediated promotion of trichome initiation in plants is conserved but utilizes species- and trichome-specific regulatory mechanisms. Plant Signaling & Behavior. 5, 205-207. (10.1104/pp.108.125385)

4 Christensen, S. A., Nemchenko, A., Borrego, E., Murray, I., Sobhy, I. S., Bosak, L., DeBlasio, S., Erb, M., Robert, C. A., Vaughn, K. A., et al. 2013 The maize lipoxygenase, ZmLOX10, mediates green leaf volatile, jasmonate and herbivore-induced plant volatile production for defense against insect attack. Plant Journal. 74, 59-73. (10.1111/tpj.12101)

5 Yan, Y., Borrego, E., V, M. 2013 Jasmonate Biosynthesis, Perception and Function in Plant Development and Stress Responses. (ed.^eds. pp.: INTECH Open Access Publisher.

6 Vlot, A. C., Dempsey, D. A., Klessig, D. F. 2009 Salicylic Acid, a multifaceted hormone to combat disease. Annual Review of Phytopathologist. 47, 177-206. (10.1146/annurev.phyto.050908.135202)

7 Aloni, R., Aloni, E., Langhans, M., Ullrich, C. I. 2006 Role of cytokinin and auxin in shaping root architecture: regulating vascular differentiation, lateral root initiation, root apical dominance and root gravitropism. Annals of Botany. 97, 883-893. (10.1093/aob/mcl027)

8 Wasternack, C., Hause, B. 2013 Jasmonates: biosynthesis, perception, signal transduction and action in plant stress response, growth and development. An update to the 2007 review in Annals of Botany. Annals of Botany. 111, 1021-1058. (10.1093/aob/mct067)

9 Davies, P. J. 2010 Plant Hormones Biosynthesis, Signal Transduction, Action! 3rd ed. NY, USA: Springer.

10 Dempsey, D. A., Vlot, A. C., Wildermuth, M. C., Klessig, D. F. 2011 Salicylic Acid biosynthesis and metabolism. Arabidopsis Book. 9, 24. (10.1199/tab.0156)

11 Zoeller, M., Stingl, N., Krischke, M., Fekete, A., Waller, F., Berger, S., Mueller, M. J. 2012 Lipid profiling of the Arabidopsis hypersensitive response reveals specific lipid peroxidation and fragmentation processes: biogenesis of pimelic and azelaic acid. Plant Physiology. 160, 365-378. (10.1104/pp.112.202846)

12 Zhao, Y. 2014 Auxin biosynthesis. Arabidopsis Book. 12, 14. (10.1199/tab.0173)

13 Widemann, E., Grausem, B., Renault, H., Pineau, E., Heinrich, C., Lugan, R., Ullmann, P., Miesch, L., Aubert, Y., Miesch, M., et al. 2015 Sequential oxidation of Jasmonoyl-Phenylalanine and Jasmonoyl-Isoleucine by multiple cytochrome P450 of the CYP94 family through newly identified aldehyde intermediates. Phytochemistry. 117, 388-399. (10.1016/j.phytochem.2015.06.027)

14 Machado, R. A., Robert, C. A., Arce, C. C., Ferrieri, A. P., Xu, S., Jimenez-Aleman, G. H., Baldwin, I. T., Erb, M. 2016 Auxin is rapidly induced by herbivory attack and regulates systemic, jasmonate-dependent defenses. Plant Physiology. (10.1104/pp.16.00940)

15 Feussner, I., Wasternack, C. 2002 The lipoxygenase pathway. Annu Rev Plant Biol. 53, 275-297. (10.1146/annurev.arplant.53.100301.135248)

16 Erb, M., Meldau, S., Howe, G. A. 2012 Role of phytohormones in insect-specific plant reactions. Trends Plant Sci. 17, 250-259. (10.1016/j.tplants.2012.01.003)

17 Chinchilla-Ramírez, M., Borrego, E. J., DeWitt, T. J., Kolomiets, M. V., Bernal, J. S. 2017 Maize seedling morphology and defence hormone profiles, but not herbivory tolerance, were mediated by domestication and modern breeding. Annals of Applied Biology. 170, 315-332. (10.1111/aab.12331)

18 Palmer, N. A., Basu, S., Heng-Moss, T., Bradshaw, J. D., Sarath, G., Louis, J. 2019 Fall armyworm (Spodoptera frugiperda Smith) feeding elicits differential defense responses in upland and lowland switchgrass. PLoS One. 14, e0218352. (10.1371/journal.pone.0218352)

19 He, Y., Borrego, E. J., Gorman, Z., Huang, P. C., Kolomiets, M. V. 2020 Relative contribution of LOX10, green leaf volatiles and JA to wound-induced local and systemic oxylipin and hormone signature in Zea mays (maize). Phytochemistry. 174, 112334. (10.1016/j.phytochem.2020.112334)

20 Borrego, E. J., Kolomiets, M. V. 2016 Synthesis and Functions of Jasmonates in Maize. Plants (Basel). 5, (10.3390/plants5040041)

21 Christensen, S. A., Nemchenko, A., Park, Y. S., Borrego, E., Huang, P. C., Schmelz, E. A., Kunze, S., Feussner, I., Yalpani, N., Meeley, R., et al. 2014 The Novel Monocot-Specific 9-lipoxygenase ZmLOX12 Is Required to Mount an Effective Jasmonate-Mediated Defense Against Fusarium Verticillioides in Maize. Molecular Plant Microbe Interaction. 27, 1263-1276. (10.1094/MPMI-06-13-0184-R)

22 Christensen, S. A., Huffaker, A., Kaplan, F., Sims, J., Ziemann, S., Doehlemann, G., Ji, L., Schmitz, R. J., Kolomiets, M. V., Alborn, H. T., et al. 2015 Maize death acids, 9-lipoxygenase-derived cyclopente(a)nones, display activity as cytotoxic phytoalexins and transcriptional mediators. Proceedings of the National Academy of Sciencie USA. 112, 11407-11412. (10.1073/pnas.1511131112)

23 Wang, K. D., Borrego, E. J., Kenerley, C. M., Kolomiets, M. V. 2020 Oxylipins Other Than Jasmonic Acid Are Xylem-Resident Signals Regulating Systemic Resistance Induced by Trichoderma virens in Maize. Plant Cell. 32, 166-185. (10.1105/tpc.19.00487)

24 Maschinski, J., Whitham, T. G. 1989 The Continuum of Plant responses to Herbivory: The Influence of Plant Asssociation, Nutrient Availability, and Timing. The American Naturalist. 134, 1–19.

25 Rowen, E., Kaplan, I. 2016 Eco-evolutionary factors drive induced plant volatiles: a meta-analysis. New Phytol. 210, 284-294. (10.1111/nph.13804)

26 Kaplan, I., Halitschke, R., Kessler, A., Sardanelli, S., Denno, R. F. 2008 Constitutive and Induced Defenses to Herbivory in Above- and Belowground Plant Tissues. Ecology. 89, 392–406.

27 Luthe, D. S., Gill, T., Zhu, L., Lopez, L., Pechanova, O., Shivaji, R., Ankala, A., Williams, W. P. 2011 Aboveground to belowground herbivore defense signaling in maize. A two-way street? Plant Signaling & Behavior. 6, 126–129.

28 Johnson, S. N., Erb, M., Hartley, S. E. 2016 Roots under attack: contrasting plant responses to below- and aboveground insect herbivory. New Phytologist. 210, 413-418. (10.1111/nph.13807)

29 Papadopoulou, G. V., van Dam, N. M. 2016 Mechanisms and ecological implications of plant-mediated interactions between belowground and aboveground insect herbivores. Ecological Research. 32, 13-26. (10.1007/s11284-016-1410-7)

30 Schmelz, E. A., Alborn, H. T., Banchio, E., Tumlinson, J. H. 2003 Quantitative relationships between induced jasmonic acid levels and volatile emission in Zea mays during Spodoptera exigua herbivory. Planta. 216, 665-673. (10.1007/s00425-002-0898-y)

31 Onkokesung, N., Galis, I., von Dahl, C. C., Matsuoka, K., Saluz, H. P., Baldwin, I. T. 2010 Jasmonic acid and ethylene modulate local responses to wounding and simulated herbivory in Nicotiana attenuata leaves. Plant Physiology. 153, 785-798. (10.1104/pp.110.156232)

32 McConn, M., Creelman, R. A., Bell, E., Mullet, J. E., Browse, J. 1997 Jasmonate is Essential for Insect Defense in Arabidopsis. Proceedings of the Nationall Acadamy of Science USA. 94, 5473–5477.

33 Erb, M., Flors, V., Karlen, D., de Lange, E., Planchamp, C., D’Alessandro, M., Turlings, T. C. J., Ton, J. 2009 Signal signature of aboveground-induced resistance upon belowground herbivory in maize. The Plant Journal. 59, 292-302. (10.1111/j.1365-313X.2009.03868.x)

34 Hasegawa, S., Sogabe, Y., Asano, T., Nakagawa, T., Nakamura, H., Kodama, H., Ohta, H., Yamaguchi, K., Mueller, M. J., Nishiuchi, T. 2011 Gene expression analysis of wounding-induced root-to-shoot communication in Arabidopsis thaliana. Plant Cell & Environment. 34, 705-716. (10.1111/j.1365-3040.2011.02274.x)

35 Ballaré, C. L. 2011 Jasmonate-induced defenses: a tale of intelligence, collaborators and rascals. Trends in Plant Science. 16, 249-257. (10.1016/j.tplants.2010.12.001)

36 Herms, D. A., Mattson, W. J. 1992 The Dilemma of Plants: To Grow or Defend. The Quarterly Review of Biology. 63,

37 Hahn, P. G., Maron, J. L. 2016 A Framework for Predicting Intraspecific Variation in Plant Defense. Trends in Ecology & Evolution. 31, 646-656. (10.1016/j.tree.2016.05.007)

38 Zust, T., Agrawal, A. A. 2017 Trade-Offs Between Plant Growth and Defense Against Insect Herbivory: An Emerging Mechanistic Synthesis. Annual Review of Plant Biology. 68, 513-534. (10.1146/annurev-arplant-042916-040856)

39 Endara, M.-J., Coley, P. D. 2011 The resource availability hypothesis revisited: a meta-analysis. Functional Ecology. 25, 389-398. (10.1111/j.1365-2435.2010.01803.x)

40 Bazzaz, F. A., Chiariello, N. R., Coley, P. D., Pitelka, L. F. 1987 Allocating Resources to Reproduction and Defense. BioScience. 37, 58–67.

41 Rosenthal, J. P., Dirzo, R. 1997 Effects of life history, domestication and agronomic selection on plant defence against insects: Evidence from maize and wild relatives. Evolutionary Ecology. 11, 337–355.

42 Troyer, A. F. 1999 Background of U.S. Hybrid Corn. Crop Science. 39, 601–626.

43 Rodriguez-Saona, C., Vorsa, N., Singh, A. P., Johnson-Cicalese, J., Szendrei, Z., Mescher, M. C., Frost, C. J. 2011 Tracing the history of plant traits under domestication in cranberries: potential consequences on anti-herbivore defences. Journal of Experimental Botany. 62, 1-12. (10.1093/jxb/erq466)

44 Whitehead, S. R., Turcotte, M. M., Poveda, K. 2017 Domestication impacts on plant-herbivore interactions: a meta-analysis. Philosophical Transactions of the Royal Society B: Biological Sciences. 372, (10.1098/rstb.2016.0034)

45 Coley, P. D., Bryant, J. P., Chapin, F. S. I. 1985 Resource Availability and Plant Antiherbivore Defense. Science. 230, 895–899.

46 Bellota, E., Medina, R. F., Bernal, J. S. 2013 Physical leaf defenses - altered by *Zea* life-history evolution, domestication, and breeding - mediate oviposition preference of a specialist leafhopper. Entomologia Experimentalis et Applicata. 149, 185-195. (10.1111/eea.12122)

47 Dávila-Flores, A. M., DeWitt, T. J., Bernal, J. S. 2013 Facilitated by nature and agriculture: performance of a specialist herbivore improves with host-plant life history evolution, domestication, and breeding. Oecologia. 173, 1425-1437. (10.1007/s00442-013-2728-2)

48 de Lange, E. S., Balmer, D., Mauch-Mani, B., Turlings, T. C. J. 2014 Insect and pathogen attack and resistance in maize and its wild ancestors, the teosintes. New Phytologist. 204, 329-341. (10.1111/nph.13005)

49 Maag, D., Erb, M., Bernal, J. S., Wolfender, J. L., Turlings, T. C., Glauser, G. 2015 Maize Domestication and Anti-Herbivore Defences: Leaf-Specific Dynamics during Early Ontogeny of Maize and Its Wild Ancestors. PLoS One. 10, e0135722. (10.1371/journal.pone.0135722)

50 Bernal, J. S., Medina, R. F. 2018 Agriculture sows pests: how crop domestication, host shifts, and agricultural intensification can create insect pests from herbivores. Current Opinion in Insect Science. 26, 76-81. (10.1016/j.cois.2018.01.008)

51 Fontes-Puebla, A. A., Bernal, J. S. 2020 Resistance and Tolerance to Root Herbivory in Maize Were Mediated by Domestication, Spread, and Breeding. Front Plant Sci. 11, 223. (10.3389/fpls.2020.00223)

52 Zou, J., Rogers, W. E., Siemann, E. 2007 Differences in morphological and physiological traits between native and invasive populations of *Sapium sebiferum*. Functional Ecology. 21, 721-730. (10.1111/j.1365-2435.2007.01298.x)

53 Chen, Y. H. 2016 Crop domestication, global human-mediated migration, and the unresolved role of geography in pest control. Elementa: Science of the Anthropocene. 4, (10.12952/journal.elementa.000106)

54 Wright, S. I., Bi, I. V., Schroeder, S. G., Yamasaki, M., Doebley, J., McMullen, M. D., Gaut, B. 2005 The Effects of Artificial Selection on the Maize Genome. Science. 308, 1310–1314.

55 Matsuoka, Y., Vigouroux, Y., Goodman, M. M., Sanchez, G. J., Buckler, E., Doebley, J. 2002 A single domestication for maize shown by multilocus microsatellite genotyping. Proceedings of The National Academy of Science USA. 99, 6080-6084. (10.1073/pnas.052125199)

56 Labate, J. A., Lamkey, K. R., Mitchell, S. E., Kresovich, S., Sullivan, H., Smith, J. S. C. 2003 Molecular and Histoical Aspects of Corn Belt Dent Diversity. Crop Science. 43,

57 Buckler, E. S. I., Stevens, N. M. 2006 Maize Origins, Domestication, and Selection. In Darwin’s Harvest, New Approaches to the Origins, Evolution, and Conservation of Crops. (ed.^eds. J. Motley, Z. Nyree, H. Cross), pp. N.Y.: Columbia University Press.

58 Hufford, M. B., Martinez-Meyer, E., Gaut, B. S., Eguiarte, L. E., Tenaillon, M. I. 2012 Inferences from the historical distribution of wild and domesticated maize provide ecological and evolutionary insight. PLoS One. 7, e47659. (10.1371/journal.pone.0047659)

59 van Heerwaarden, J., Hufford, M. B., Ross-Ibarra, J. 2012 Historical genomics of North American maize. Proceedings of the National Academy of Science USA. 109, 12420-12425. (10.1073/pnas.1209275109)

60 Cecchini, N. M., Roychoudhry, S., Speed, D. J., Steffes, K., Tambe, A., Zodrow, K., Konstantinoff, K., Jung, H. W., Engle, N. L., Tschaplinski, T. J., et al. 2019 Underground Azelaic Acid-Conferred Resistance to Pseudomonas syringae in Arabidopsis. Molecular Plant-Microbe Interactions. 32, 86-94. (10.1094/MPMI-07-18-0185-R)

61 Zhao, Y. 2010 Auxin biosynthesis and its role in plant development. Annual Review of Plant Biology. 61, 49-64. (10.1146/annurev-arplant-042809-112308)

62 Robert, C. A., Veyrat, N., Glauser, G., Marti, G., Doyen, G. R., Villard, N., Gaillard, M. D., Köllner, T. G., Giron, D., Body, M., et al. 2012 A specialist root herbivore exploits defensive metabolites to locate nutritious tissues. Ecology Letters. 15, 55-64. (10.1111/j.1461-0248.2011.01708.x)

63 Abdi, H. 2007 The Bonferonni and Sidak Corrections for Multiple Comparisons. In Encyclopedia of Measurement and Statistics. (ed.^eds. N. Salkind), pp. Thousand Oaks, CA: SAGE.

64 Richardson, J. T. E. 2011 Eta squared and partial eta squared as measures of effect size in educational research. Educational Research Review. 6, 135-147. (10.1016/j.edurev.2010.12.001)

65 Yeo, I. K., Johnson, R. A. 2000 A new family of power transformations to improve normality or symmetry. Biometrika. 87, 954–959.

66 SAS Institute Inc. JMP. Pro 14.0.0 ed. Cary, NC 2018.

67 Lai, X., Yan, L., Lu, Y., Schnable, J. C. 2018 Largely unlinked gene sets targeted by selection for domestication syndrome phenotypes in maize and sorghum. Plant J. 93, 843-855. (10.1111/tpj.13806)

68 Castano-Duque, L., Helms, A., Ali, J. G., Luthe, D. 2018 Plant Bio-Wars: Maize Protein Networks Reveal Tissue-Specific Defense Strategies in Response to a Root Herbivore. Chemical Ecology. 44, 727-745. (10.1007/s10886-018-0972-y)

69 Agrawal, A. A., Salminen, J. P., Fishbein, M. 2009 Phylogenetic trends in phenolic metabolism of milkweeds (Asclepias): evidence for escalation. Evolution. 63, 663-673. (10.1111/j.1558-5646.2008.00573.x)

70 Cipollini, D., Walters, D., Voelckel, C. 2014 Costs of Resistance in Plants: From Theory to Evidence. In Annual Plant Reviews online. (ed.^eds. pp. 263–307

71 Ruiz, C. J. A., Sánchez, G. J., Aguilar, S. M. 2001 Potential Geographical Distribution of Teosinte in Mexico: A GIS Approach. Maydica. 46, 105–110.

72 Sanchez Gonzalez, J. J., Ruiz Corral, J. A., Garcia, G. M., Ojeda, G. R., Larios, L. C., Holland, J. B., Medrano, R. M., Garcia Romero, G. E. 2018 Ecogeography of teosinte. PLoS One. 13, e0192676. (10.1371/journal.pone.0192676)

73 Bellon, M. R., Taylor, J. E. 1993 “Folk” Soil Taxonomy and the Partial Adoption of New Seed Varieties. Economic Development and Cultural Change. 41, 763-786. (10.1086/452047)

74 Bellon, M. R., Mastretta-Yanes, A., Ponce-Mendoza, A., Ortiz-Santamaria, D., Oliveros-Galindo, O., Perales, H., Acevedo, F., Sarukhan, J. 2018 Evolutionary and food supply implications of ongoing maize domestication by Mexican campesinos. Proceedings of the Royal Society B. 285, (10.1098/rspb.2018.1049)

75 Bellon, M. R., Hodson, D., Bergvinson, D., Beck, D., Martinez-Romero, E., Montoya, Y. 2005 Targeting agricultural research to benefit poor farmers: Relating poverty mapping to maize environments in Mexico. Food Policy. 30, 476-492. (10.1016/j.foodpol.2005.09.003)

76 Bellon, M. R., Adato, M., Becerril, J., Mindek, D. 2003 The Impact of Improved Maize Germplasm on Poverty Alleviation: The Case of Tuxpeño-derived Material in Mexico. International Food Policy Research Institue.

77 Orozco–Ramírez, Q., Brush, S. B., Grote, M. N., Perales, H. 2014 A Minor Role for Environmental Adaptation in Local–Scale Maize Landrace Distribution: Results from a Common Garden Experiment in Oaxaca, Mexico1. Economic Botany. 68, 383-396. (10.1007/s12231-014-9285-4)

78 Perkins, J. H. 1982 Insects, Experts, and the Insecticide Crisis: The Quest for New Pest Management Strategies. Plenum Press.

79 Kutka, F. 2011 Open-Pollinated vs. Hybrid Maize Cultivars. Sustainability. 3, 1531-1554. (10.3390/su3091531)

80 Duvick, D. N. 2005 The Contribution of Breeding to Yield Advances in maize (*Zea mays* L.). In Advances in Agronomy (ed.^eds. pp. 83–145

81 Stintzi, A., Weber, H., Reymond, P., Browse, J., Farmer, E. E. 2001 Plant defense in the ansence of jasmonic acid: The role of cyclopentenones. Proceedings of The National Academy of Science USA. 98, 12837–12842.

82 Varsani, S., Grover, S., Zhou, S., Koch, K. G., Huang, P. C., Kolomiets, M. V., Williams, W. P., Heng-Moss, T., Sarath, G., Luthe, D. S., et al. 2019 12-Oxo-Phytodienoic Acid Acts as a Regulator of Maize Defense against Corn Leaf Aphid. Plant Physiology. 179, 1402-1415. (10.1104/pp.18.01472)

83 Fonseca, S., Chini, A., Hamberg, M., Adie, B., Porzel, A., Kramell, R., Miersch, O., Wasternack, C., Solano, R. 2009 (+)-7-iso-Jasmonoyl-L-isoleucine is the endogenous bioactive jasmonate. Nature Chemical Biology. 5, 344-350. (10.1038/nchembio.161)

84 Heitz, T., Widemann, E., Lugan, R., Miesch, L., Ullmann, P., Desaubry, L., Holder, E., Grausem, B., Kandel, S., Miesch, M., et al. 2012 Cytochromes P450 CYP94C1 and CYP94B3 catalyze two successive oxidation steps of plant hormone Jasmonoyl-isoleucine for catabolic turnover. J Biol Chem. 287, 6296-6306. (10.1074/jbc.M111.316364)

85 Baldwin, I. T. 1998 Jasmonate-induced responses are costly but benefit plants under attack in native populations. Proc. Natl. Acad. Sci. 95, 8113–8118.

86 Diezel, C., Allmann, S., Baldwin, I. T. 2011 Mechanisms of optimal defense patterns in Nicotiana attenuata: flowering attenuates herbivory-elicited ethylene and jasmonate signaling. J Integr Plant Biol. 53, 971-983. (10.1111/j.1744-7909.2011.01086.x)

87 Poudel, A. N., Zhang, T., Kwasniewski, M., Nakabayashi, R., Kazuki, S., Koo, A. J. 2016 Mutations in jasmonyl-L-isoleucine-12-hydroxylases suppress multiple JA-dependent wound responses in *Arabidopsis thaliana*. Biochimica et Biphysica Acta. 1861, 1396–1408. (https://doi-org.srv-proxy2.library.tamu.edu/10.1016/j.bbalip.2016.03.006)

88 Park, S. W., Li, W., Viehhauser, A., He, B., Kim, S., Nilsson, A. K., Andersson, M. X., Kittle, J. D., Ambavaram, M. M., Luan, S., et al. 2013 Cyclophilin 20-3 relays a 12-oxo-phytodienoic acid signal during stress responsive regulation of cellular redox homeostasis. Proceedings of The National Academy of Science USA. 110, 9559-9564. (10.1073/pnas.1218872110)

89 Hazman, M., Suhnel, M., Schafer, S., Zumsteg, J., Lesot, A., Beltran, F., Marquis, V., Herrgott, L., Miesch, L., Riemann, M., et al. 2019 Characterization of Jasmonoyl-Isoleucine (JA-Ile) Hormonal Catabolic Pathways in Rice upon Wounding and Salt Stress. Rice (N Y). 12, 45. (10.1186/s12284-019-0303-0)

90 Gao, X., Starr, J., Gobel, C., Engelberth, J., Feussner, I., Tumlinson, J., Kolomiets, M. 2008 Maize 9-Lipoxygenase ZMLOX3 Controls Development, Root-Specific Expression of Defense Genes, and Resistance to Root-Knot Nematodes. Molecular Plant-Microbe Interactions. 21, 98–109.

91 Hwang, I. S., Hwang, B. K. 2010 The pepper 9-lipoxygenase gene CaLOX1 functions in defense and cell death responses to microbial pathogens. Plant Physiology. 152, 948-967. (10.1104/pp.109.147827)

92 Jung, H. W., Tschaplinski, T. J., Wang, L., Glazebrook, J., T., G. J. 2009 Priming in Systemic Plant Immunity. Science. 324, 89–91.

93 Gao, Q. M., Zhu, S., Kachroo, P., Kachroo, A. 2015 Signal regulators of systemic acquired resistance. Frontiers in Plant Science. 6, 228. (10.3389/fpls.2015.00228)

94 Wu, L., Wang, S., Chen, X., Wang, X., Wu, L., Zu, X., Chen, Y. 2013 Proteomic and phytohormone analysis of the response of maize (Zea mays L.) seedlings to sugarcane mosaic virus. PLoS One. 8, e70295. (10.1371/journal.pone.0070295)

95 Gaudin, A. C. M., McClymont, S. A., Raizada, M. N. 2011 The Nitrogen Adaptation Strategy of the Wild Teosinte Ancestor of Modern Maize, subsp. Crop Science. 51, 2780. (10.2135/cropsci2010.12.0686)

96 Gaudin, A. C. M., McClymont, S. A., Soliman, S. S. M., Raizada, M. N. 2014 The effect of altered dosage of a mutant allele of *Teosinte branched 1 (tb1-ref)* on the root system of modern maize. Genetics. 15,

97 Gao, X., Shim, W.-B., Gobel, C., Kunze, S., Feussner, I., Meeley, R., Balint-Kurtim, P., Kolomiets, M. 2007 Disruption of Maize 9-Lipoxygenase Results in Increased Resistance to Fungal Pathogens and Reduced Levels of Contamination with Mycotoxin Fumonisin. Molecular Plant-Microbe Interactions. 20, 922–933.

98 Chen, Q., Sun, J., Zhai, Q., Zhou, W., Qi, L., Xu, L., Wang, B., Chen, R., Jiang, H., Qi, J., et al. 2011 The basic helix-loop-helix transcription factor MYC2 directly represses PLETHORA expression during jasmonate-mediated modulation of the root stem cell niche in Arabidopsis. Plant Cell. 23, 3335-3352. (10.1105/tpc.111.089870)

99 Sun, J., Chen, Q., Qi, L., Jiang, H., Li, S., Xu, Y., Liu, F., Zhou, W., Pan, J., Li, X., et al. 2011 Jasmonate modulates endocytosis and plasma membrane accumulation of the Arabidopsis PIN2 protein. New Phytol. 191, 360-375. (10.1111/j.1469-8137.2011.03713.x)

100 Blossey, B., Notzold, R. 1995 Evolution of Increased Competitive Ability in Invasive Nonindigenous Plants: A Hypothesis. British Ecological Society. 83, 887–889.

101 Erb, M., Balmer, D., De Lange, E. S., Von Merey, G., Planchamp, C., Robert, C. A., Roder, G., Sobhy, I., Zwahlen, C., Mauch-Mani, B., et al. 2011 Synergies and trade-offs between insect and pathogen resistance in maize leaves and roots. Plant Cell & Environment. 34, 1088-1103. (10.1111/j.1365-3040.2011.02307.x)

102 Gray, M. E., Sappington, T. W., Miller, N. J., Moeser, J., Bohn, M. O. 2009 Adaptation and invasiveness of western corn rootworm: intensifying research on a worsening pest. Annual Review of Entomology. 54, 303-321. (10.1146/annurev.ento.54.110807.090434)

103 Lombaert, E., Ciosi, M., Miller, N. J., Sappington, T. W., Blin, A., Guillemaud, T. 2017 Colonization history of the western corn rootworm (*Diabrotica virgifera virgifera*) in North America: insights from random forest ABC using microsatellite data. Biological Invasions. (10.1007/s10530-017-1566-2)

104 Baker, H. G. 1972 Human Influences on Plant Evolution. Economic Botany. 26, 32–43.

105 McMullen, M. D., Frey, M., Degenhardt, J. 2009 Genetics and Biochemistry of Insect Resistance in Maize. In Handbook of Maize: Its Biology. (ed.^eds. pp. 271–289

